# Targeting the sphingolipid rheostat in IDH1*^mut^* glioma alters cholesterol homeostasis and triggers apoptosis via membrane degradation

**DOI:** 10.1101/2024.04.26.591321

**Authors:** Tyrone Dowdy, Helena Muley Vilamu, Adrian Lita, Aiguo Li, Tomohiro Yamasaki, Lumin Zhang, Raj Chari, Hua Song, Meili Zhang, Wei Zhang, Nicole Briceno, Dionne Davis, Mark R. Gilbert, Mioara Larion

**Affiliations:** Neuro-Oncology Branch, National Cancer Institutes, National Institutes of Health, Bethesda, Maryland; Genome Modification Core Laboratory, National Cancer Institutes, National Institutes of Health, Frederick, Maryland

**Keywords:** N, N-dimethylsphingosine, sphingosine, ceramide, IDH1*^mut^* gliomas, sphingolipid, reverse cholesterol transport, HMGCS, HMGCR, ABCA1, ABCG1, MYLIP, IDOL, ACAT2, LXR nuclear receptor, oligodendroglioma, astrocytoma, lipidomics, multi-omics

## Abstract

The cross-regulation of metabolism and trafficking is not well understood for the vital sphingolipids and cholesterol constituents of cellular compartments. While reports are starting to surface on how sphingolipids like sphingomyelin (SM) dysregulate cholesterol levels in different cellular compartments (Jiang et al., 2022), limited research is available on the mechanisms driving the relationship between sphingolipids and cholesterol homeostasis, or its biological implications. Previously, we have identified sphingolipid metabolism as a unique vulnerability for IDH1^mut^ gliomas via a rational drug design. Herein, we show how modulating sphingolipid levels affects cholesterol homeostasis in brain tumors. However, we unexpectedly discovered for the first time that C17 sphingosine and NDMS addition to cancer cells alters cholesterol homeostasis by impacting its cellular synthesis, uptake, and efflux leading to a net decrease in cholesterol levels and inducing apoptosis. Our results reflect a reverse correlation between the levels of sphingosines, NDMS, and unesterified, free cholesterol in the cells. We show that increasing sphingosine and NDMS (a sphingosine analog) levels alter not only the trafficking of cholesterol between membranes but also the efflux and synthesis of cholesterol. We also demonstrate that despite the effort to remove free cholesterol by ABCA1-mediated efflux or by suppressing machinery for the influx (LDLR) and biosynthetic pathway (HMGCR), apoptosis is inevitable for IDH1^mut^ glioma cells. This is the first study that shows how altering sphingosine levels directly affects cholesterol homeostasis in cancer cells and can be used to manipulate this relationship to induce apoptosis in IDH1^mut^ gliomas.

## INTRODUCTION

Cholesterol is a precursor of steroid hormones, bile acids, and vitamins that are essential for cellular function, signaling, and survival (Zhao & Dahlman-Wright, 2010). The membrane cholesterol content directly impacts the biosensors that mediate the activity and expression of a cascade of membrane proteins (Morales et al., 2023; Zhao et al., 2010). Of which, low-density lipoprotein receptor (LDLR) enables the docking of apolipoprotein E (Apo E), which participates in the trafficking of LDL-cholesterols (Morales et al., 2023; Trinh et al., 2020). Following LDLR-mediated internalization, LDL-cholesterol enters the cellular space by endocytosis (Morales, 2023). During this process, early and late endosomes (EE and LE, respectively) experience the fate of homotypic fusion with lysosomes to form transient endo-lysosome organelles that known to traffic LDL-cholesterol as well as sphingolipids (Appelqvist et al., 2011; Huotari et al., 2011). Niemann–Pick C1 (NPC1) and Niemann–Pick C2 (NPC2) facilitate the export of unesterified cholesterol along with unsaturated sphingosine from the lysosome (Altuzar et al., 2023; Trinh et al., 2020). Unesterified cholesterol from the cytosol enters the endoplasmic reticulum (ER) via the assistance of the GRAM domain containing 1C (GRAMD1) (Altuzar et al., 2023; Trinh et al., 2020). After migration through the ER, biosensor feedback such as liver X nuclear receptors (LXR) from the neighboring nuclear membrane is responsible for activating the signaling pathways that mediate pro-cell survival responses and the regulation of cholesterol homeostasis (Trinh et al., 2020; Zhao & Dahlman-Wright, 2010).

Cholesterol biosensors such as LXRα (NR1H3) mainly mediate the expression and activity of membrane proteins that supervise the cholesterol export, internalization, and biosynthesis (Zelcer et al., 2009; Zhao & Dahlman-Wright, 2010; Zhu et al., 2012). Of these proteins, ATP-binding cassette transporters ABCA1 and ABCG1 moderate the export of cholesterol from the plasma membrane (PM) to extracellular ApoA1 and high-density lipoprotein (HDL) in addition to ABCG1-mediated export from ER to attenuate membrane stress (Rong et al., 2013; Tarling et al., 2011; Yvan-Charvet et al., 2010; Zhu et al., 2012). In addition, biosensors such as LXRα can indirectly suppress cholesterol trafficking by activating PM-bound myosin regulatory light chain interacting protein (MYLIP)—an E3 ubiquitin ligase also referred to as inducible degrader of the LDLR (IDOL)—activity and associated ubiquitin-proteosome degradation of LDLR upon detection of excessive cholesterol accumulation (Adi et al., 2019; Zelcer et al., 2009).

Cholesterol levels can suspend the expression of catalytic enzymes responsible for isoprenoid precursor, sterol, and cholesterol biosynthesis, which is initialized by hydroxymethylglutaryl-CoA synthase (HGMCS) activity. In relation, HMGCS forms intermediate HMG-CoA by condensing acetyl-CoA and acetoacetyl-CoA, typically supplied by β-oxidation of fatty acids ((Chen et al., 2022). The rate-limiting NADH-dependent HMG-CoA reductase (HMGCR) catalyzes the conversion of HMG-CoA to mevalonate, the essential precursor for isoprenoid, sterol, and cholesterol biosynthesis (DeBose-Boyd, 2008; H. yan Wang et al., 2022; Yeh et al., 2018). Cholesterol and sterol feedback can also trigger the binding of LXR-mediated Insulin-induced gene 1 (Insig-1) to HMGCR which recruits gp78 ubiquitin ligase to accelerate ubiquitination and proteolytic degradation of HMGCR in the ER (DeBose-Boyd, 2008; Feramisco et al., 2004; Sever et al., 2003). Interestingly, HMGCS and HMGCR have also become a recent focus in the development of anti-cancer combination treatments for their crucial role in tumor progression (Chen et al., 2022; H. yan Wang et al., 2022; Zhang et al., 2019).

Sphingolipids, including ceramide (Cer), sphingosine (Sph), sphingomyelin (SM), and sphingosine-1-phosphate (S1P), are not just bioactive molecules, but they play a crucial role in influencing vital signaling pathways (Dowdy et al., 2020). Sphingolipid-dependent signaling is a complex process that modulates cellular regulation, metabolism, proliferation, metastasis, and cell death (Dowdy et al., 2020). Herein, we show how modulating sphingolipid levels affects cholesterol homeostasis in brain tumors.

However, we unexpectedly discovered for the first time that C17 sphingosine and NDMS addition to IDH1^mut^ cells alters cholesterol homeostasis by impacting its cellular synthesis, uptake, and efflux leading to a net decrease in cholesterol levels and inducing apoptosis. Our results reflect a reverse correlation between the levels of sphingosines, NDMS, and unesterified, free cholesterol in the cells. We show that increasing sphingosine and NDMS (a sphingosine analog) levels alter not only the trafficking of cholesterol between membranes but also the efflux and synthesis of cholesterol. We also demonstrate that despite the effort to remove free cholesterol by ABCA1-mediated efflux or by suppressing machinery for the influx (LDLR) and biosynthetic pathway (HMGCR), apoptosis is inevitable for IDH1^mut^ glioma cells. This is the first study that shows how altering sphingosine levels directly affects cholesterol homeostasis in cancer cells and can be used to manipulate this relationship to induce apoptosis in IDH1^mut^ gliomas.

## METHODS

Unless specified, LC/MS grade solvents and HPLC grade eluent additives acquired from Thermo Fisher Scientific Inc. (Waltham, MA) were used to prepare extracts and mobile phases. All LC/MS and MS/MS data were acquired using high-resolution (HR) Agilent 6545 Quadrupole Time-of-Flight (QToF) Mass Spectrometer (MS) coupled with Infinity II 1290 Liquid Chromatography Ultra-High-Pressure system (Agilent Technologies Inc., Santa Clara, CA).

### Cell Culture and Sample Collection

Suspension media for human-derived cells (BT142+R132H, TS603+R132H, U251±R132H overexpression, GSC923, and Normal Human Astrocytes, NHA) were cultured using 500 mL DMEM: F12 Ham media (Gibco Laboratories, Gaithersburg, MD) with the following additives: 5 mL penicillin/streptomycin 100X; 5 mL N_2_ supplement 100X; 100 µL Epidermal growth factor (100 µg/mL EGF); 100 µL fibroblast growth factors (100 µg/mL FGF) obtained from ThermoFisher Scientific (Waltham, MA); 0.5 mL 2.0 mg/mL Heparin Sulfate (MilliporeSigma, St. Louis, MO, USA). The human-derived NCH1681 glioma cell media was prepared using 500 mL DMEM: F12/Glutamax media with the following additives: 5 mL Penicillin/Streptomycin 100X; 100 µL at 100 µg/mL EGF and FGF and 10 mL BIT Admixture 100X (PELOBiotech GmbH, Planegg-Martinsried, Germany) per 40 mL of prepared media. Following the addition of additives, each media preparation was filter-sterilized and stored at 4°C.

Neurosphere was grown at 37°C to a minimum density of 3 million cells/replicate and ∼90% cell viability per flask for RNA sequencing, lipidomic, and Western blot protein experiments. Neurospheres were partitioned into treatment groups based on DMSO control, timepoint, and drug concentration. Each treatment group was administered either sphingosine C17:1 (SphC17), N,N-dimethylsphingosine (NDMS), or a combination for RNA sequencing analysis. Based on the outcome of the RNA sequencing experiment, additional lipidomic and western blot analyses were performed to examine the drug-dependent response and to assess the mechanisms of drug action for the combination treatment only. This design was based on the RNAseq and cell viability data (Dowdy, 2020), which revealed that the combination treatment provided the most significant response and differential expression of vital genes in IDH1^mut^ gliomas.

Samples were aspirated from each flask and transferred to 15-mL pre-sterilized conical tubes and centrifuged at 400 x g for 5 min at room temperature (RT). Cell pellets were washed with 600 µL PBS and centrifuged at 400 rpm for 3 min at RT. The supernatant was discarded, and pellets were snap-frozen on dry ice and stored at −80 °C to ensure complete quenching of all metabolic activity and degradation until extraction.

### Lipid extraction

Before sonication, each sample was administered 300 µL ice-chilled MilliQ water and homogenized via probe sonication by Misonix XL-2000 Ultra-liquid processor (Misonix Inc., Farmingdale, NY, USA) at 40 amps for 30 s. Following sonication, an aliquot (5 % total volume) of each cell lysate was collected and stored at −80 °C to conduct Bradford protein quantification for later normalize each analyte measurement to sample-specific protein. During the extraction procedure, 600 µL chilled (−20 °C) LC/MS grade methanol (MeOH) was added to each sample lysate. Each sample was vortexed for 15 s and placed on an Orbi-blotter mixing rotator (Benchmark Scientific, Edison, NJ, USA) at max speed to incubate in an ice bath for 10 min. Each sample was administered 500 µL chilled (−20 °C) chloroform followed by 120 µL HPLC grade chloroform IS solution containing 0.05 µg/mL 3-phenyl-N-(4-pyridinyl)-acrylamide (PNPA). After each sample was vortexed for 1 min, the samples were returned to the ice bath and placed on a rotator for 30 min to separate phases using this adapted Bligh-Dyer multi-phase extraction adapted from Dowdy et al. (2020). Samples were centrifuged at 13000 x g for 20 min at 4°C. The lipid extract was transferred from the resulting multiple phases (upper hydrophilic, lower hydrophobic lipid, and intermediate protein disk). All samples remained in an ice-bath until final centrifugation and collection. The retained lipid extracts were concentrated on Techne sample concentrator with PTFE-coated needles (Cole-Palmer, Vernon Hills, IL) under high purity (grade 4.5) N_2_ gas (Robert Oxygen, Gaithersburg, MD) flow until completely dry, snap frozen, and then stored at −80 °C.

### Biostatic Drug Dose-response Analysis

The biostatic effects of the drug treatment were investigated using the CCK-8 cell viability assay (Dojindo Molecular Technologies, Inc. Rockville, MD) as previously described (Dowdy,2020.). The media (50 μl) prepared at the desired drug concentration or equivalent DMSO (v/v%) for control were added to designated wells of each Corning sterile 96-well non-treated black chimney, transparent round bottom plate for spheroids (Sigma-Aldrich, Inc., St. Louis, MO). Then, 50 μl cell suspension media was added to each well after being diluted based on a prior optimization experiment (Dowdy et al., 2020). Plates were covered with a lid and placed in a cell CO_2_ incubator at 37℃. CCK-8 indicator solution (10 μl) was added to each well of the plate before measuring the absorbance at 515 nm using a microplate reader. The blanks were composed of media and 0.3% DMSO to subtract the background from each well measurement. The growth response was determined based on the percent of absorbance per well to the absorbance of associated DMSO control (containing untreated cells) from the same row. The mean response per treatment group was reported as percent growth per DMSO control.

### Enriched sphingolipid, sterol, and polar lipid LC/MS detection

Multiple IDH1^mut^ neurospheres (oligodendroglioma TS603, oligoastrocytoma BT142 and NCH1681, genetically overexpressed IDH1^mut^ model U251-R132H) and representative IDH1^wt^ neurospheres (GSC923 and U251^wt^) were prepared as described (section 4.1). Treated groups were administered 6.5 µM SphC17 combined with 6.5 µM NDMS, an SphK1 inhibitor (SphK1i), to promote the accumulation of sphingosines and ceramides from the De Novo Sphingolipid Biosynthesis Pathway. Prior to analysis, hydrophobic lipid extracts were reconstituted in 120 µL 5:3:1 Molecular Biology Grade Ethanol (EtOH)/ MeOH/ water, vortexed for 3 min, centrifuged (13000 x g, 4 °C) for 5 min, and transferred to amber glass LC vial with 0.250 mL deactivated glass insert and caps with PTFE/ silicone septa (Agilent Technologies Inc.). The pooled quality control (QC) sample was prepared using 10% volume from each sample. As described by Dowdy et al., lipids were resolved using the coupled column LC method that was optimized to enhance retention and resolution of sphingolipids, particularly as well as sterols and other polar lipids by incorporating Xbridge BEH amide 2.5 µm, 2.1 x 50 mm column (Waters Corp) coupled with a pentafluorophenyl (PFP) InfinityLab Poroshell 120, 1.9 µm, 2.1 x 100 mm (Agilent Technologies Inc.). The enrichment of sphingolipids and polar lipids over a gradient composed of mobile phase A, 30 mM ammonium acetate (NH_4_Ac) (aq) + 0.1% formic acid (FA) + 0.0015% InfinityLab Deactivator (Agilent Technologies, Inc.) in 1% MeOH, pH 4.3; mobile phase B, 5:10:85 30mM NH_4_Ac/MeOH/ACN, pH 4.3. An isothermal column temperature of 45°C and a 0.200 mL/min flow rate were applied along with a gradient timetable: 0-1.5 min, 2 % B; 3.5 min, 67% B, 5.5 min 100 % B; hold 1.5 min; 8 min, 75 % B; 8.5 min, 100% B; hold 0.75 min; 11.5 min, 2 % B, equilibrate, 1.5 min. Real-time mass correction was applied with 0.2 mL/min infusion of external standard (containing TFA/HP921) in 95:5 ACN/water. Dynamic range electrospray injection (ESI) negative ion acquisition applied the following MS parameters: injection volume, 7.5 µL drying gas temperature (temp), 250°C; gas flow 9 L/min; nebulizer, 45 psi; sheath gas temp, 350°C; sheath gas flow, 12 L/min; capillary voltage, 3000 V; nozzle voltage, 15 V; fragmentor, 90 V; skimmer, 50 V; scan rate, 3.0 spectra/sec; mass range 75-1300 m/z. Alternatively, ESI positive ion acquisition applied the following MS parameters: injection volume, 6.5 µL drying gas temperature (temp), 250°C; gas flow 9 L/min; nebulizer, 45 psig; sheath gas temp, 350°C; sheath gas flow, 12 L/min; capillary voltage, 3500 V; nozzle voltage, 15 V; fragmentor, 125 V; skimmer, 50 V; scan rate, 3.0 spectra/sec; mass range 75-1300 m/z.

### Global lipidomic LC/MS analysis

To ensure broad coverage for all lipid classes of interest (mainly sphingolipids and sterols), global lipidomic LC/MS analysis was also performed using Acquity UPLC CSH 1.7 µm, 2.1 × 100 mm column (Waters Corp. Milford, MA, USA) using the LC gradient mobile phase A—30% ACN (aq), 5 mM Ammonium Formate and 0.15% FA—and mobile phase B—90:10 Isopropanol (IPA)/ACN, 5 mM Ammonium Formate, and 0.15% FA—while applying the LC and MS conditions previously described for global lipidomic analysis ((Larion et al., 2019)).

### Time-dependent, relative quantification of cholesterol

Representative IDH1^mut^ neurospheres (TS603) and associated media were prepared as described in the section. Treated groups were administered 6.5 µM SphC17 combined with 6.5 µM NDMS as described above Section 4.6.2). Prior to analysis, cell and media (200 µL) lipid extracts were reconstituted in 120 µL 5:3:1 EtOH/MeOH/water, vortexed for 3 min, centrifuged (13000 x g, 4 °C) for 5 min, and transferred to amber glass LC vial with 0.250 mL deactivated glass. The pooled QC sample was prepared using 10% volume of each sample extract. Cholesterol was detected using LC method incorporating Xbridge BEH amide 2.5 µm, 2.1 x 50 mm column coupled with a PFP InfinityLab Poroshell 120, 1.9 µm, 2.1 x 100 mm; over a gradient composed of mobile phase A—30 mM NH_4_Ac (aq) + 0.1% FA+ 0.0015% InfinityLab Deactivator in 1%MeOH (aq), pH 4.3—and mobile phase B—5:10:85 30mM NH_4_Ac/MeOH/ACN, pH 4.3. A flow rate of 0.200 mL/min was applied along with a gradient timetable: initial, 1% B; 0.5 min, 2% B; 3.5 min, 67% B, 5.5 min, 100 % B; hold 4 min; 9.6 min, 70 % B; 11 min, 95% B; 11.5 min, 100 % B; hold 1.0 min; equilibrate, 1.5 min. To reduce carryover, the following column temperature gradient was applied during the wash phase of each run: initial, 50°C until 8.5 min; 9-13 min, 60°C; 13.5 min, 50°C. To maintain the carryover ≤ 20% with intra-day and inter-day (2 d) CV%≤ 20% acceptable for bioanalytical assays ((Clouser-Roche et al., 2008), it was necessary to perform intermediate a post-run wash injection following each standard and sample injection. An isothermal 60°C column temperature was applied for the post-run wash gradient composed of mobile phase A—30 mM NH_4_Ac + 0.1% FA+ 0.0015% InfinityLab Deactivator in 1% MeOH (aq), pH 4.3—and mobile phase B— 95:5ACN/MeOH + 0.2% FA. A flow rate of 0.200 mL/min was applied along with a gradient timetable: initial, 99% B; 4 min, 100% B; 5.5 min, 67% B; 6.5 min, 95% A; 7.5 min, 100% B; 8.2 min, 99% B; 9.2% B, 40% B; 10.2 min, 1% B; equilibrate, 1.0 min at 50°C column temperature.

The acquisition MS methods included real-time mass correction applied with 0.2 mL/min infusion of external standard (containing TFA/HP921) in 95:5 ACN/water. High-resolution ESI positive acquisition applied the following parameters: injection volume, 6.0 µL; drying gas temp, 250°C; gas flow 9 L/min; nebulizer, 45 psig; sheath gas temp, 350°C; sheath gas flow, 12 L/min; capillary voltage, 3500 V; nozzle voltage, 15 V; fragmentor, 125 V; skimmer, 50 V; scan rate, 3.0 spectra/sec; mass range 118-1700 m/z. In addition, an MS/MS method was applied to a commercial grade standard, pooled QC, and spiked QC sample using collision-induced dissociation at 12V, targeted MS/MS 3.0 spectra/sec scan rate to validate that the target ion for cholesterol (m/z 369.3516; transition, m/z 215.1) [M-H2O+H]^+^ was detected at retention time (RT) of 7.0 min. A standard curve ranging from 0.03 µg/mL to 13.0 µg/mL (unweighted, linear R^2^ = 0.99) was prepared using the ratio of Cholesterol/ internal standard to measure the relative concentration of cholesterol per sample.

### LC/MS data processing and analysis

Prior to preprocessing each dataset, pooled QC samples (TIC, BPI, and EIC) were chromatographically examined to inspect the consistency of retention time and ionization levels throughout. Following acquisition, mass feature bins were defined by partitioning the m/z vs. RT matrices into fixed width using Agilent Masshunter Profinder B.08.00. Bins were manually inspected to confirm consistent, reproducible integration for each compound of interest across all samples. The logical binning for each neutral mass assignment was determined using molecular feature extraction algorithm within Profinder to deconvolute, integrate, and envelope parent ions, selected adducts (H- and H+ only), and natural isotopes to define each composite spectrum based on the precursor ion. In Profinder, targeted ion selection, alignment, binning, and annotation for each compound was restricted to mass accuracy ±2.0 mDa, precursor ion abundance ≥ 1000, and retention time ± 0.5 min using a proprietary, gradient-specific Masshunter Personal Compound Data Library (PCDL) as a reference for pre-determined compound ion targets. Following pre-processing, the ion area for each analyte was adjusted to the ratio by area of the sample-specific internal standard and then normalized based on sample-specific Bradford protein (µg) quantification in Microsoft Excel to generate csv file. The normalized values from csv file were uploaded and log-transformed in Metaboanalyst 5.0 software (Pang et al., 2021) to conduct multivariate analysis (Pearson-Ward correlation heatmap, principal component analysis, and volcano plot using non-parametric, Mann-Whitney U t-test) for each group comparison. Graphs were also generated using GraphPad Prism 10.0, applying non-parametric, Mann-Whitney U t-test and representing related p-value as: p ≤ 0.05, *; p ≤ 0.005, **; p ≤ 0.0005, **; p ≤ 0.00005, ****; not significant, ns.

### RNA purification

The PureLink Mini Kit (Thermo Fisher Scientific) was used to extract and purify RNA from IDH1^mut^ glioma neurospheres to assess transcriptomic similarities as a function of treatment. Neurospheres were grown until they reached 3 million/replicate and treated SphC17, NDMS, or the combination and collected as described above(Section 4.1). The extraction and purification were conducted as previously described in the PureLink® RNA Mini Kit User Guide (https://www.thermofisher.com/document-connect/document-connect.html?url= https://assets.thermofisher.com%2FTFS-Assets%2FLSG%2Fmanuals%2Fpurelink_rna_mini_kit_man.pdf&title=UHVyZUxpbmsgUk5BIE1pbmkgS2l0). The following molecular biology grade materials were also used to complete the RNA purification procedure: PureLink DNase (Thermo Fisher Scientific); DNase-, RNase- and Protease-Free 200 proof Ethanol ≥99.5%, BioUltra 2–mercaptoethanol and RNAase/Nuclease-free Water (Sigma-Aldrich). Upon completion of the purification, the RNA extracts were quantified using nanodrop, aliquoted, snap frozen, and then stored at −80 °C. Samples were shipped on dry ice to the NIH Center for Cancer Research CCR Sequencing Core Facility (CCR-SF, Frederick, MD) for global detection, identification, and relative quantification of the transcriptome.

### RNAseq data processing and analysis

Sequencing reads of RNA-seq raw data were analyzed using CCBR Pipeliner y9hi(https://github.com/CCBR/Pipeliner). The pipeline performs several tasks: pre-alignment reads quality control, grooming of sequencing reads, alignment to the human reference genome (hg38), post-alignment reads quality control, feature quantification, and differentially-expressed gene identification. In the QC phase, the sequencing quality of each sample is independently assessed using FastQC, Preseq, Picard tools, RSeQC, SAMtools, and QualiMap. Sequencing reads that passed quality control thresholds were trimmed for adaptor sequences using the Cutadapt algorithm. The transcripts were annotated and quantified using STAR. Sample quality control was vigorously carried out based on sequencing criteria such as mapping rates, library complexities, etc. In cases where differentially-expressed genes were derived, further sample filtering was performed based on the zero value of specific genes and the sample variance. Differentially-expressed genes were examined using LIMMA and EdgeR statistical methodology to determine DEGs with the thresholds of p < 0.05 and |FC| =1.3 based on best fit. Graphs were also generated using GraphPad Prism 10.0, applying non-parametric, Mann-Whitney U t-test and representing related p-value as: p ≤ 0.05, *; p ≤ 0.005, **; p ≤ 0.0005, **; p ≤ 0.00005, ****; not significant, ns.

### Immunoblot protein analysis

Cellular lysates from representative IDH1mut TS603 and IDH1wt GSC923 neurospheres treated with drug combination (6.5 µM SphC17 and 6.5 µM NMDS) or DMSO (control) were prepared using NP-40 lysis buffer containing protease inhibitors (Thermo Fisher Scientific Inc.). Lysates were centrifuged at 13,000 × g for 10 min to obtain the supernatants. Pierce BCA protein assay (Thermo Fisher Scientific Inc.) was performed for protein quantification. The proteins in the lysates were resolved by SDS-polyacrylamide gel electrophoresis (BIO-RAD, Hercules, CA) and were transferred to Low-fluorescence (LF) PVDF membranes (BIO-RAD). Membranes were blotted with Tris Buffered Saline with Tween®20 (TBS-T) buffer (Cell Signaling Technology) containing 5% of nonfat milk (BIO-RAD). TBS-T buffer was used for all the membrane washings. Primary antibodies were diluted in TBS-T buffer with 0.2% Bovine Serum Albumin (BSA) (Roche). Membranes were incubated with the primary antibody dilution at 4°C overnight. After washings, membranes were incubated with HRP-conjugated secondary antibody (Jackson IR) at room temperature for 1 hour. The bands were visualized using an enhanced chemiluminescence (ECL) system (ChemiDoc MP, BIO-RAD). Equal loading of proteins was verified by immunoblotting for β-actin or α-tubulin. The primary antibodies were mainly acquired from AB (Abcam Inc., Waltham, MA) or CST (Cell Signaling Technology) as follows: Mouse anti-ABCA1 (1:1000, AB66217), rabbit anti-ABCG1 (1:1000, AB52617), rabbit anti-SREBP1 (1:200, AB3259), rabbit anti-LDL receptor (1:500, AB52818), mouse anti-HMGCR (1:1000, AB242315), rabbit anti-HMGCS1 (1:1000, CST36877), rabbit anti-cleaved PARP1 (1:1000, AB32064), rabbit anti-Cytochrome c (1:1000, CST11940), anti-β-actin (1:5000, AB6276), mouse anti-α-tubulin (1:1000; CST3873) and mouse anti-IDH1 (R132H) (1:500, SAB4200548, Sigma-Aldrich).

### Fluorescence microscopy

Lice cell microscopy images were collected using a Leica STELLARIS 8 CRS (Coherent Raman Scattering) microscope equipped with FALCON and fluorescence lifetime imaging (FLIM) and using a 40x water objective HC PL APO 10x/0.40 CS2 NA 0.4. The detectors used were HyDS1 and HyDX. Hoersch was imaged using excitation at 350 nm and emission around 462, Bodipy-Cholesterol (Ex: 480nm, Em: 508nm); Rhodamine-C17 sphingosine (Ex: 555nm, Em: 580nm) and plasma membrane (Ex:645nm, Em: 666nm).

## RESULTS

Herein we examined the mechanisms of drug action and cell death for IDH1^mut^ gliomas that were treated with Sph C17, NDMS, or the combination. We elucidated a unique relationship between sphingolipids (e.g., sphingosine) and their astonishing ability to interact and trigger the regulation of sterols (e.g., cholesterol). Here, we reported on the events triggered by a negative correlation between sphingosines and C18 sphingosine-derived NDMS along with cholesterol. This negative correlation was directly connected to cytosolic accumulation of sphingosines (and its derivatives), their ability to activate expression for genes involved in LXR-mediated signaling pathways, and the direct impact leading to the biostatic response across IDH1^mut^ oligodendroglioma and astrocytoma subtypes.

Initially, the cytotoxic dose-dependent response that occurred across multiple IDH1^mut^ glioma (figure 1b-c) subtypes Sph C17 and NDMS (figure 1a) was attributed to an accumulation of presumed apoptotic sphingolipids (i.e., sphingosine and ceramides) while simultaneously inhibiting the catalytic conversion of sphingosine to sphingosine-1-phosphate (S1P) by SphK1. This assumption was based on earlier reports that IDH1^mut^ gliomas presented profoundly lower global levels of sphingolipids (including sphingosine and its derivatives) compared to IDH1 wild-type GBM (Dowdy et al., 2020). Accordingly, the time-dependent increase of CYT C and cPARP1 proteins (associated with membrane-stress-driven cell death) in representative IDH1^mut^ neurospheres surpassed representative IDH1^wt^ neurospheres (figure 1b). This phenotypic distinction provided evidence to further validation for a potential metabolic vulnerability that could be manipulated by targeting IDH1^mut^ gliomas using Sph and NDMS. As expected, multiple IDH1^mut^ glioma subtypes consistently demonstrated enhanced sensitivity to combination treatment (figure 1c) when compared to IDH^1^ wild-type glioblastoma (GSC923) as well as normal human astrocyte (NHA) neurospheres.

**Figure 1.**
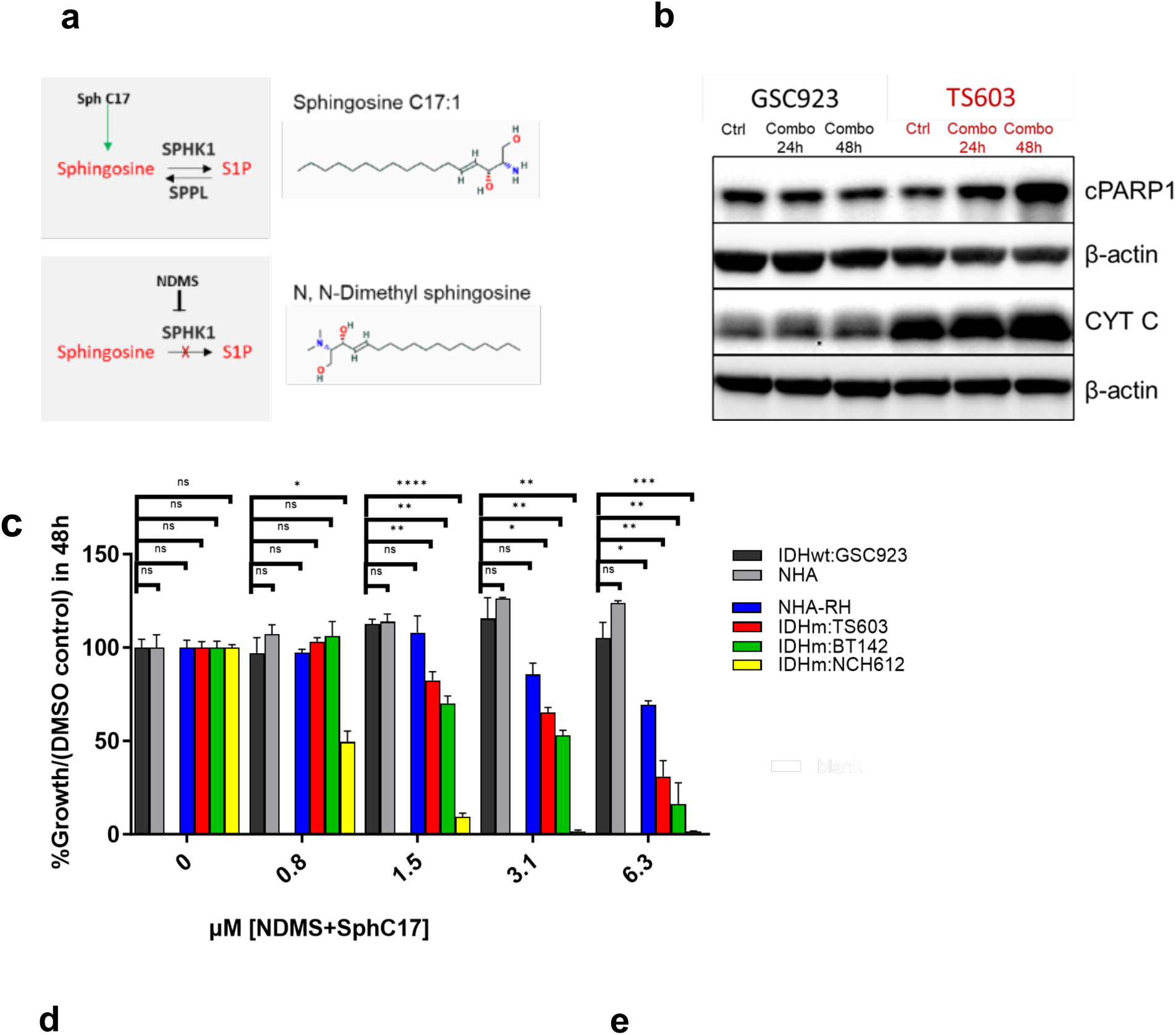
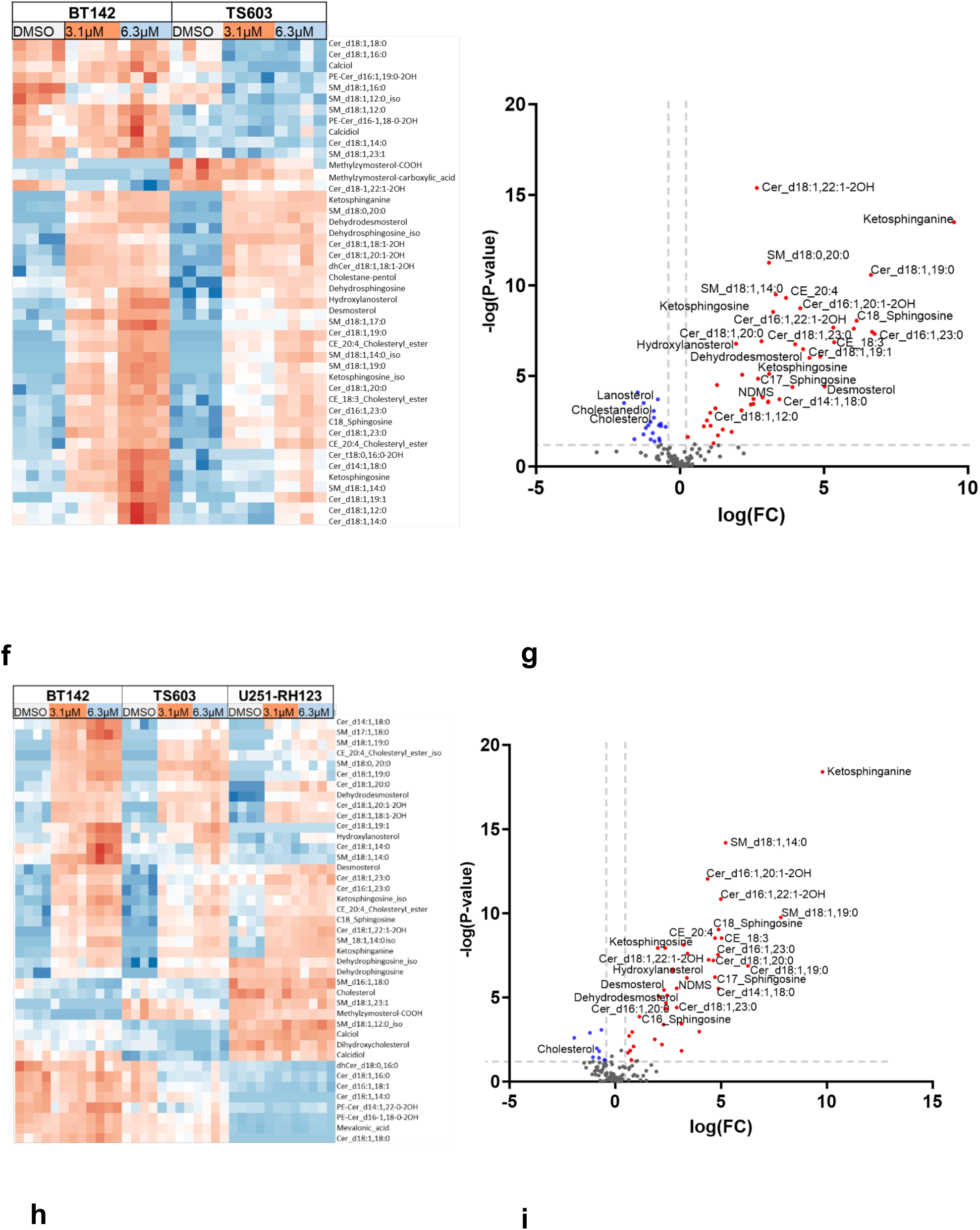
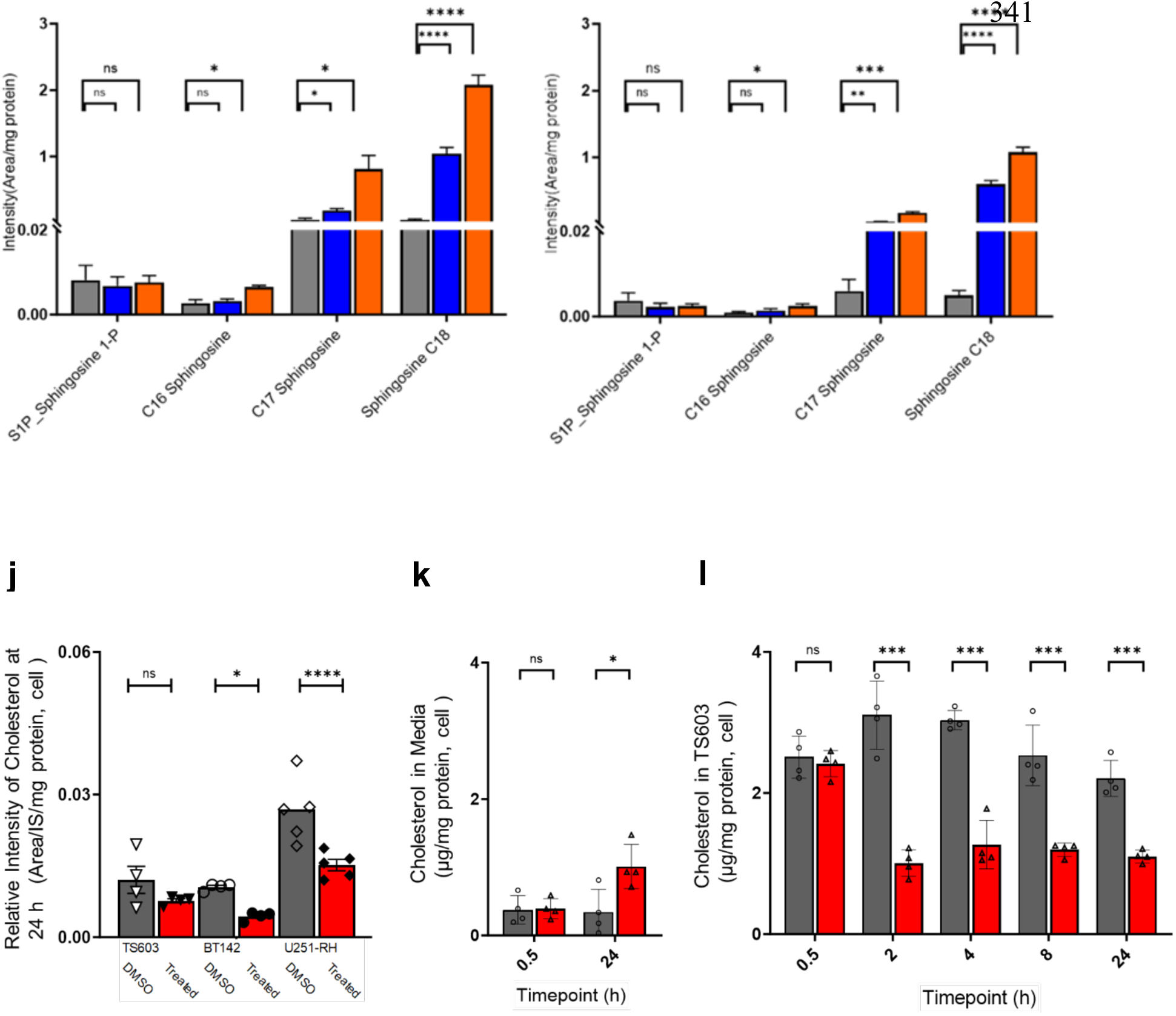
LC/MS analysis with sphingolipids and sterols enrichment using combination treatment with Sph C17 and SphK1 inhibitor NDMS after 1 d. **(a)** The dose-dependent biostatic effect that Sph C17 and sphingosine kinase (SphK) inhibitor, NDMS induced in highly susceptible IDH1^mut^ glioma subtypes was attributed to a subsequent accumulation of pro-apoptotic sphingosine while suppressing the formation S1P by with SphK1 activity. **(b)** Immunoblot (n = 3) demonstrated an apparent elevation in the apoptotic markers, cytochrome C (CYT C), and cleaved PARP1 (cPARP1) that was outstanding for representative IDH1^mut^ TS603 neurospheres compared to the representative IDH1^wt^ GSC923 neurospheres following treatment with an elevated concentration at 6.5 µM SphC17 and NDMS. **(c)** CCK8 cell assay demonstrated an enhanced susceptibility to the combination treatment for representative IDH1^mut^ oligodendroglioma (TS603) and astrocytomas (BT142 and NCH612) compared to glioblastoma (GSC923) as well as normal human astrocyte (NHA) neurospheres. The NHA that were genetically modified to overexpressed mutant IDH1-R132H (NHA-RH) also presented an elevated sensitivity to combination treatment. **(d)** The unsupervised hierarchical correlation heatmap illustrated the global dysregulation of sphingolipids and sterols across the representative IDH1^mut^ oligodendroglioma TS603 and astrocytoma BT142. **(e)** The consequent volcano plot (cut-off FC=|1.3|; p-value ≤ 0.05) demonstrates that significant elevation of sphingosine bases (sph) and particular sphingolipids (certain sphingomyelins, ceramides, and oxidized ceramides) is negatively correlated to the downregulation of cholesterol and certain derivative products. **(f)** A complementary heatmap demonstrated the reproducibility of these trends across representative, nascent IDH1^mut^ gliomas along with the model U251 with genetic overexpression of the IDH1-R132H mutation. **(g)** The consequent volcano plot (cut-off FC=|1.3|; p-value≤ 0.05) reveals that the elevation of all sph (C16; C17; C18) bases along with certain sphingolipids were conserved along with their outstanding negative correlation to cholesterol. While heatmaps **(d, f)** and volcano plots **(e, g)** are only able to display one feasible putative IDs for acylated features, a repository table is available with comprehensive details on the m/z, adducts, retention time, and alternate ID nomenclature. Legend: CE, cholesterol ester (cholesteryl); Cer, ceramide; dhCer, dihydroceramide; PE-Cer, ceramide phosphoethanolamine; Sph, sphingosine; SM, sphingomyelin; S1P, sphingosine-1-phosphate. **(h)** Comparative analysis for all sphingosine bases confirmed that their accumulation as a function of drug combination treatment (6.25 µM, red; 3.125 µM, blue; DMSO control, grey) as anticipated in representative astrocytoma (BT142) and **(i)** conserved in oligodendroglioma (TS603). Additionally, the observed decline in S1P did not appear to be significant, suggesting that it is not the main drive for phenotypic changes and related biostatic effects of the treatment in IDH1^mut^ gliomas. **(j)** Comparison of the relative levels provides evidence that the decline in cholesterol is conserved across nascent and genetically modified IDH1^mut^ neurospheres following 24h combination treatment (6.25 µM, red; DMSO control, grey). **(k)** A time-course LC/MS experiment (inter-/intra-day CV% ≤ 20%; carryover ≤ 20%; linear curve R^2^ = 0.99) provided a further assessment of the relative concentration of cholesterol released into the media at the initial (0.5 h) and final (24 h) timepoints following treatment with 6.25 µM combination (red) or DMSO (grey) in the representative IDH1^mut^ neurospheres. **(l)** The time-course LC/MS experiment was extended (at 0.5, 2, 4, 8, and 24 h) to also probe the unexpected decline in relative cholesterol levels (figure 1j) as a function drug treatment (6.25 µM, red; DMSO control, grey). Bar graphs display the mean per group, error bars represent the standard error of the mean (SEM), and nonparametric, Mann-Whitney U t-test p-values are defined as: *, p ≤ 0.05; **, p ≤ 0.005; ***, p ≤ 0.0005; ****, p ≤ 0.00005; ns, not significant.

The Pearson-Ward correlation heatmap (figure 1d)—which only compared response in IDH1^mut^ oligodendroglioma and astrocytoma neurospheres to assess common trends across IDH1^mut^ subtypes— and a related volcano plot (figure e) revealed that certain sphingolipids (i.e. sphingosines, ceramides, and sphingomyelins), sterols (e.g. desmosterol and other oxysterols), and CE formed clusters that are positively correlated across IDH1^mut^ glioma following combination treatment. A Pearson-Ward correlation heatmap (figure 1f) was prepared to under which trends were conserved upon analyzing neurospheres from IDH1^mut^ oligodendroglioma and astrocytoma, and GBM with genetic overexpression of IDH1^mut^ neurospheres after 24 h treatment with drug combination. The multivariate analysis with the heatmap (figure 1d and 1f) and volcano plots (figure 1e and 1g) presented a profoundly consistent trend involving the global elevation of sphingosines (sph) and other non-phosphorylated sphingolipids alongside certain cholesterol precursors in response to combination drug.

Moreover, the significant elevation of certain C18:1 and C17:1 sphingomyelin (SM) (figure 1d-g) was also consistent with the elevation in Sph C18 and Sph C17 (figure 1d-g). The sphingosines (sph C16, C17, and C18) detected were all consistently elevated (figure d-i) across multiple IDH1^mut^ glioma neurospheres during 24 h treatment with NDMS and Sph C17 combination. Interestingly, cholesterol levels declined (figure 1d-g and 1j) across multiple IDH1^mut^ neurospheres despite an elevation in certain sterol precursors and CEs that clustered with sphingosine and related derivatives. In order to determine whether the observed trends resulted from early-on, drug-dependent elevation in cholesterol biosynthesis, the relative cholesterol concentrations were measured over time (figure k-l). Time course measurement of cholesterol concentrations (figure 1l) in TS603 treated (red) with DMSO control (gray) as well as the media (figure 1k).

The assessment of the relative concentration of cholesterol released in media (figure 1k) at the initial (0.5 h) and final (24 h) time points suggested that the efflux of cholesterol drove the phenotypic metabolic change in the representative IDH1^mut^ neurospheres. Relative cholesterol levels decline similarly during the first 24 h across multiple IDH1-mutant neurospheres (figure 1d-g and 1l). The time-course LC/MS experiment probed the unexpected alteration in the relative concentration of cholesterol (figure 1l) as a function of drug treatment and the resultant metabolic change accumulation of sph bases along with certain other sphingolipids (figure d-i). The data (figure 1k-l) suggested that basal cholesterol levels (figure 1j) were not recovered after reaching a plateau within 2 h after treatment. Unexpectedly, an early onset surge in cholesterol concentration was not observed. Such an event could have accounted for the activation of the LXR-related cholesterol homeostasis mechanism, which was prevalent throughout the transcriptomic data and will be discussed further.

### Transcriptome revealed sphingosine alters cholesterol sensing pathway

We performed an initial pilot transcriptomic study (Figure 2a-b) with representative IDH1^mut^ astrocytoma to capture early onset response to targeting sphingolipid metabolism of IDH1^mut^ gliomas by enriching intracellular sphingosine level (fig 1d-i). Herein IDH1^mut^ neurospheres were treated for 24 h at concentration of the combination with NDMS and Sph C17 combination that presented reduced toxicity compared to higher dose. SGMS2 (figure 2a) was among the only sphingolipid enzymes to be significantly elevated in the transcriptome for representative IDH1^mut^ (figure 2a) following the 24 h combination treatment. It is possible that the neurospheres shunted sphingosine towards sphingomyelinase (SGMS), given that we found the drug combination treatment with Sph C17 along with NDMS SphKi elevated sphingosines (figure 1h-i) and sphingomyelin. In accordance, the significant elevation of certain C18:1 and C17:1 sphingomyelin (SM) complemented the elevation in both C18 and C17 sphingosine (figure 1d-g).

**Figure 2.**
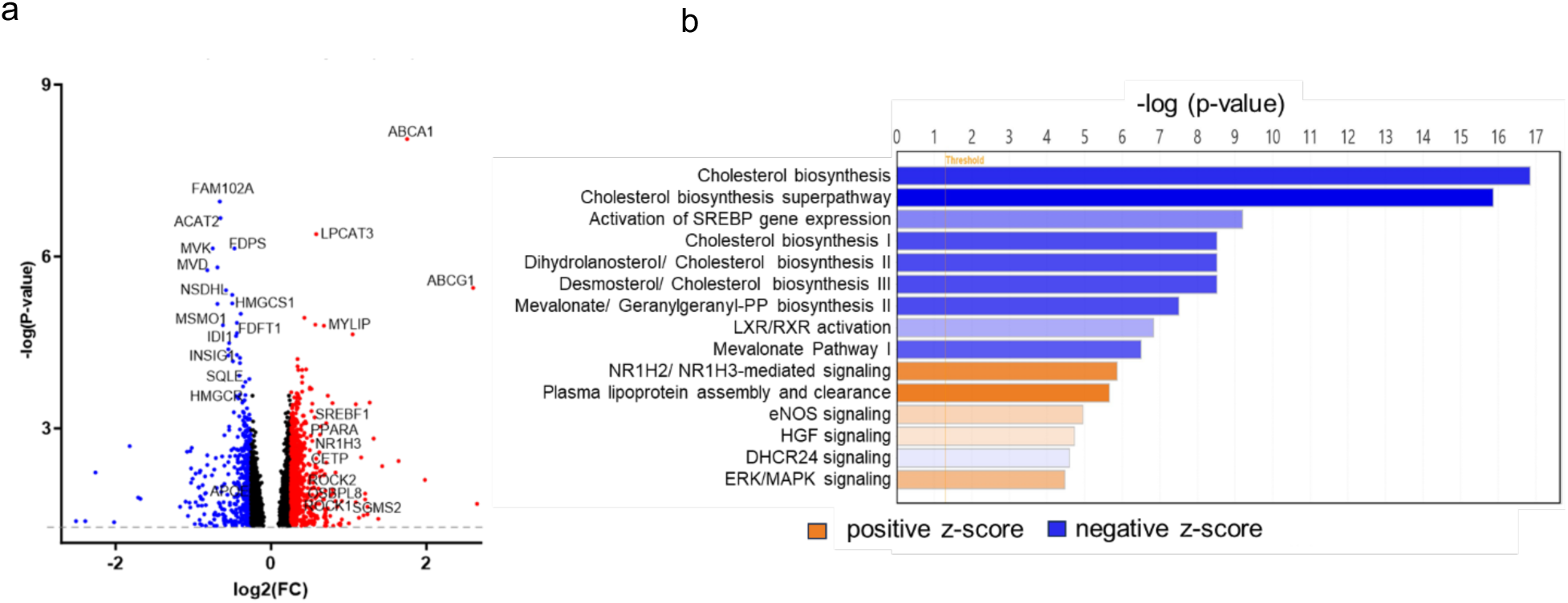
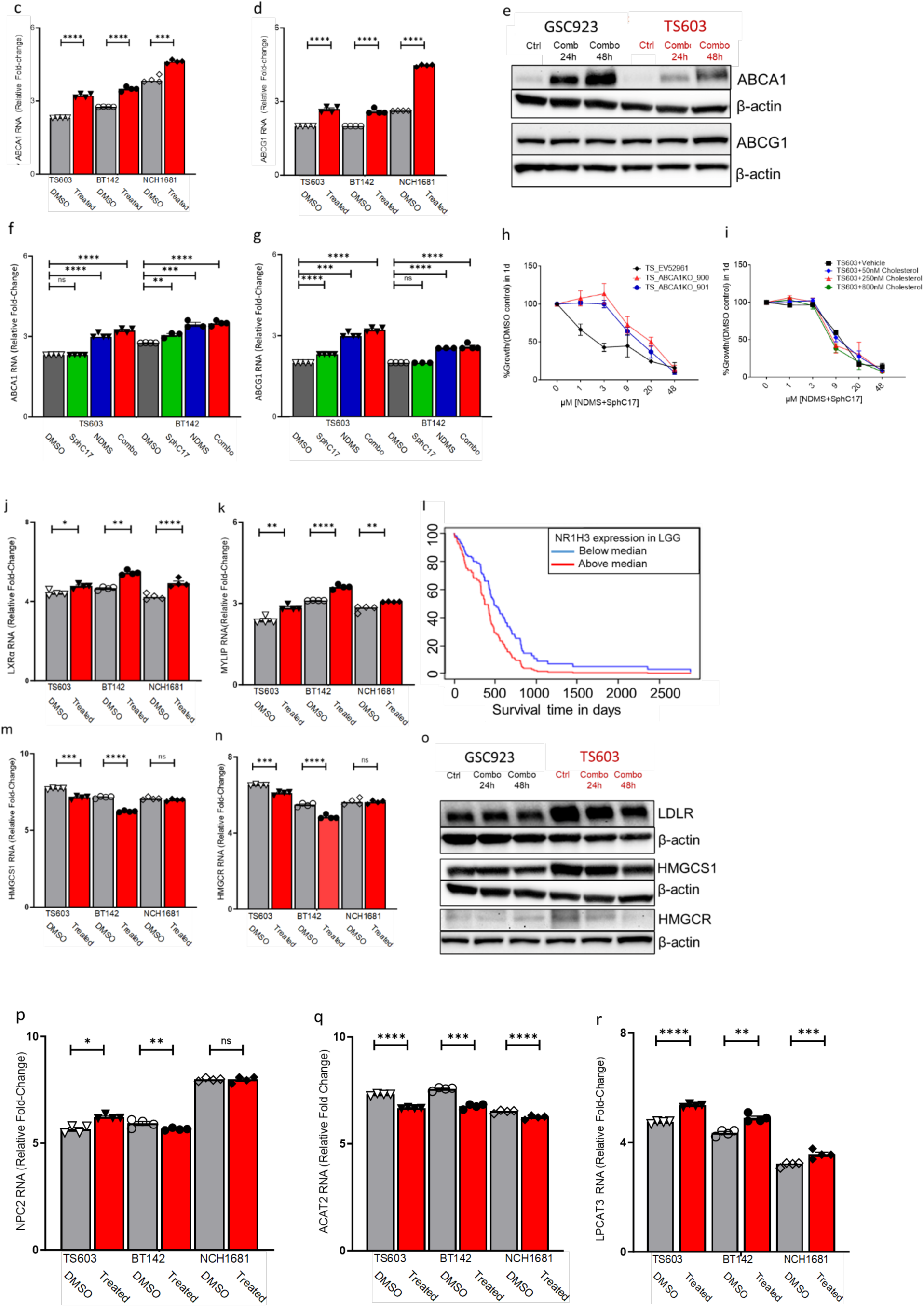
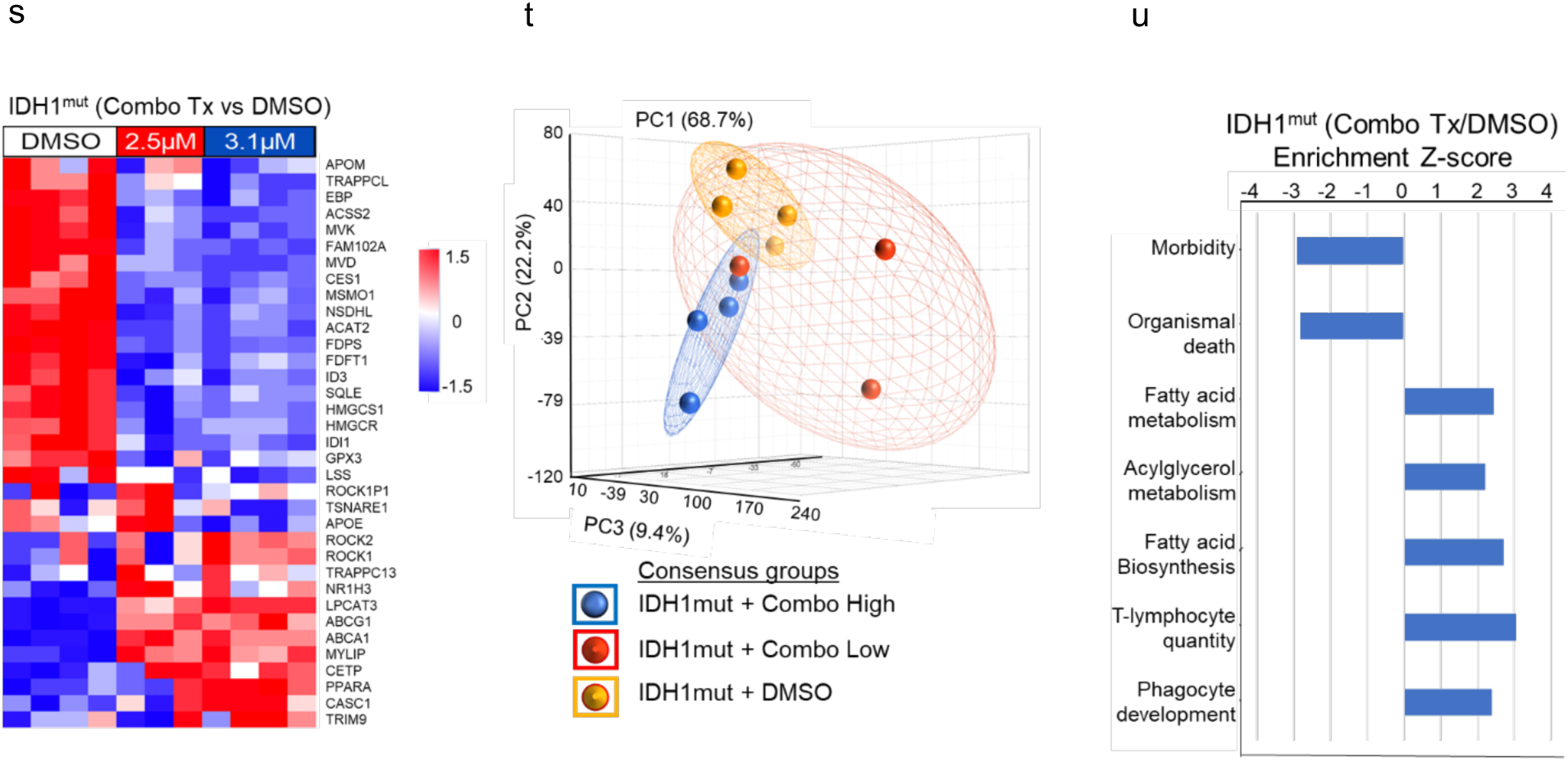
RNA and Protein biomarker expression was determined for IDH1^mut^ glioma spheroids treated for 24 and 48 h with Sph C17, NDMS, or the combination (Combo). **(a)** The volcano plot (cutoff FC ≈ |1.3|; p-value ≤ 0.05; n = 3) illustrates that the most prominent expression involves genes associated with the LXR signaling pathway, mainly genes for proteins that participate in controlling intracellular free and membrane-bound cholesterol by facilitating the efflux (ABCA1, ABCG1 and LPCAT3) of cholesterol primarily, ubiquitinated proteolytic degradation of LDLR influx machinery (MYLIP), and LXRα (NR1H3) are elevated. In contrast, the labeled downregulated genes (particularly HMGCR and HMGCS1) are highly significant and all related to the biosynthesis of cholesterol, other sterols, and their isoprenoid precursors following 24h treatment at a low concentration (3.25 µM Combo) to investigate early-onset changes in representative IDH1^mut^ neurospheres (BT142)**. (b)** Related Ingenuity pathway analysis (FC ≈ |1.3|; p ≤ 0.05) found that 33 genes involved in mevalonate and cholesterol biosynthesis were suppressed and negatively correlated with the elevation of 123 genes related to the activation of LXR/RXR. **(c)** (ABCA1 and **(d)** ABCG1 levels were significantly elevated within transcriptome for broad coverage of IDH1^mut^ neurospheres including various glioma subtypes (oligodendroglioma TS603 and astrocytomas BT142 and NCH1681) following 48h following 48h at 6.5 µM NDMS + SphC17 combo treatment to emphasize expression at a critical point when a higher apoptotic marker was evident through immunoblot analysis for a less sensitive, representative IDH1^mut^ (TS603) compared to representative IDH1^wt^ (GSC923) neurospheres(figure 1b) and all IDH1^mut^ glioma subtypes would experience cellular cycle arrest (figure 1c). **(e)** The immunoblot blot reveals proteomic elevation of ABCA1 and ABCG1 in representative IDH1^mut^ and IDH1^wt^ neurospheres in response to 6.5 µM Combo treatment after 48 h. The transcriptomic expression of d) ABCA1 and e) ABCG1 confirms a drug-dependent response across multiple subtypes of IDH1^mut^ neurospheres (TS603, BT142, and NCH1681) after 48 h at 6.5 µM Combo treatment. **(f)** RNA levels of ABCA1 and **(g)** ABCG1 appear to be the most elevated NDMS and the combination of NDMS with Sph C17 when compared to Sph C17 alone. **(h)** The graph (n = 3) for 24 h Combo treatment dose-response study with CCK8 revealed that the representative IDH1^mut^ neurospheres ΔABCA1 knockout 900 vectors and ΔABCA1 knockout 901 vectors presented higher tolerance for Combo treatment when compared to control empty vector EV 52961. **(i)** Unexpectedly, the graph (n = 3) 24 h Combo treatment dose-response in representative IDH1^mut^ neurospheres did not appear to be rescued or otherwise impacted by 2 h pre-incubation at various cholesterol (0, 50, 250 and 800 nM) concentrations. **(j)** The graph provides evidence that even within 48 h, the activation and expression of LXRα remained significantly elevated across a range of IDH1^mut^ glioma subtypes in response to the Combo treatment. **(k)** Graph confirms that gene expression of LXR-mediated MYLIP was significant at 48 h across multiple IDH1^mut^ glioma subtypes and was consistent with **(a)** early-on MYLIP expression at 24 h. **(l)** The Kaplan-Meier survival curve comparison with Cox proportional-hazards age model expression (hazard ratio = 1.71; p-value =0.02) was generated using The Glioblastoma Bio Discovery Portal (GBM-BioDP) resource to access The Cancer Genome Atlas (TCGA) data for clinical patients with low-grade gliomas (LGG) subclass c(Celiku et al., 2014). **(m)** The graph confirmed that HMGCS1 and **(n)** HMGCR transcription declined significantly across multiple IDH1^mut^ glioma subtypes during the 48 h Combo treatment as expected downstream of LXRα (NR1H3) activation and signaling. **(o)** Immunoblot analysis presented a complementary decline in the protein levels for the LDLR (the primary target of MYLIP-driven proteolytic ubiquitination), and the HMGCS1 and HMGCR1 (the rate-limiting onset for the biosynthesis of cholesterol, and its precursors) observed across IDH1^mut^ gliomas subtypes in response to 48 h Combo treatment. **(p)** Under the same conditions, a decline was observed for the transcription of NPC2 the export of unesterified from the lysosome **(q)** Likewise, a decline was conserved for the transcription of ACAT2 across IDH1^mut^ gliomas subtypes after 48 h Combo treatment. **(r)** An elevation in the transcription for LXR-mediated LPCAT3 was apparent after 48 h treatment. **(s)** Hierarchical clustering of the most differentiated genes revealed dose-dependent expression across the three treatment conditions. **(t)** Principal component analysis based on the dysregulation of all RNA from neurosphere samples from three treatment conditions. Ellipsoids represent standard deviation. **(u)** Enrichment analysis based on dysregulation of the transcriptome.

While the prominent transcription markers did not provide direct insight into common markers of cell death (figure 1b), the volcano plot (figure 2a) and hierarchical clustering (figure 2s) revealed an interesting elevation in the transcription of Rho-associated coiled-coil containing protein kinase 1 (ROCK1), which is known to mediate plasma membrane stress-driven vesiculation (also apoptotic membrane blebbing)(Tixeira et al., 2020). The volcano plot (figure 2a) and hierarchical clustering (figure 2s) mainly highlighted a stark contrast in the expression between the downregulation in the transcription for gene involved in catalyzing the biosynthesis of cholesterol and its precursors (FDPS, HMGCS, HMGCR, IDI1, MVD, MVK, NSDHL, SQLE) and concomitant upregulation in the transcription of genes (NR1H3 also known as LXRα, ABCA1, ABCG1, LPCAT3, CETP and MYLIP) involved in the LXR-related signaling to reduce excess intracellular, unesterified cholesterol as a consequence of drug treatment.

Additionally, the Ingenuity pathway enrichment analysis (Figure 2b) demonstrated a profound downregulation in the transcription for a broad cluster of genes involved in sterol biosynthesis pathways such as the Mevalonate, Cholesterol, Farnesyl pyrophosphate (Farnesyl-PP), Zymosterol pathways. The apparent global decline included the transcription of genes for proteins involving the mevalonate (MVA) pathway associated with isoprenoid biosynthesis upstream of Sterol and Cholesterol biosynthesis.

Herein, the overall suppression of RNA and protein expression for a broad range of catalytic enzymes involved in MVA, Sterol, and Cholesterol pathway, including rate-limiting steps (HMGCS1, HMGCR, and LSS), was attributed to the sphingosine-rich NDMS and Sph C17 treatment combination (figure 2a-b, and 2m-o). In previous studies, the expression of catalytic enzymes (HMGCR and lanosterol synthase, LSS) involved in sterol and Cholesterol biosynthesis was positively attributed to GBM malignancy (Phillips et al., 2019; Qiu et al., 2016). Moreover, HMGCR has also become the focus point in tumor progression research due to its pivotal function in the production of precursors for cholesterol and other sterol biosynthesis (Chen et al., 2022; Qiu et al., 2016; H. yan Wang et al., 2022; Zhang et al., 2019).

The observed downregulation in the transcription of genes involved in cholesterol and related precursor biosynthesis is negatively correlated with the upregulation in the transcription of genes involved in LXR-mediated signaling to regulate cholesterol homeostasis (figure 2a-g, 2j-k, and 2m-o). While the activation of LXR-mediated signaling has been typically attributed to the LXR biosensor response to the accumulation of lipotoxic cholesterol, our findings revealed a rapid decline, rather than an excess, in intracellular cholesterol over the time-course experiment (figure 1j). The time-course experiment (figure 1k) also confirmed that cholesterol efflux accounted for a portion of the intracellular loss, given that cholesterol content in the media increased substantially by the 24 h time point.

We conducted further investigations to probe dysregulation in gene transcription that was enhanced or preserved across a broader selection of IDH1^mut^ glioma subtypes (categorized as astrocytoma or oligodendroglioma) over the extended 48 h period at an elevated concentration of the NDMS, Sph C17 or the combination treatment. Given that most of the cells had undergone cell cycle arrest (figure 1b), fewer of the changes that were evident at the early onset pilot study remained significant upon a follow- on at 48 h treatment. However, the primary trend was preserved, thus revealing the decline in the transcription of genes for catalytic enzymes involved in the biosynthesis of cholesterol and its precursors (e.g., HMGCS1 and HMGCR) and concomitant elevation in the transcription of genes that participate in LXR-mediated cholesterol efflux (particularly, ABCA1 and ABCG1) remained the most significant changes across as function of drug treatment. Moreover, the preserved significance in the elevation in the transcription of ABCA1 and ABCG1 across all tested IDH1^mut^ glioma subtypes concurs with our interpretation that cholesterol efflux was a major contributing to the observed cellular decline and subsequent elevation of cholesterol in the media.

Furthermore, both sphingosine C18-derived NDMS and Sph C17 promoted the expression of the gene for ABCA1 and ABCG1 transporters of the reverse cholesterol transport pathway independently across multiple IDH1^mut^ subtypes (figure 2f-g). In relation, C18 sphingosine was also significantly elevated along with the other sphingosines detected in the lipidome following treatment (figure 1e and 1g) across multiple IDH1^mut^ subtypes.

Interestingly, Transcriptomic analysis revealed that the gene expression for the LXRα (NR1H3) was significantly elevated in the first 24 h and 48 h following treatment IDH1^mut^ neurospheres compared to untreated control. It has been previously reported that the LXR target ABCG1 can accelerate the efflux of cholesterol from ER to reduce ER-stress related apoptosis in response to cytotoxic accumulation of cholesterol (Alnaaim et al., 2024; Röhrl et al., 2014; T. Wang et al., 2020). These findings also supported the hypothesis that the activation of LXRα and related transcription of downstream targets (i.e., ABCA1, ABCG1, MYLIP) that regulate cholesterol homeostasis in conjunction with the global suppression of genes associated with cholesterol biosynthesis is part of LXR-related biosensor rescue response to protect cells from a perceived cytotoxic accumulation of cholesterol in the cellular membrane.

Given that the immunoblot revealed that representative IDH1^wt^ GSC923 (figure 2e) presented greater elevation of ABCA1 in response to treatment, these representative IDH1^wt^ GBM may have additional adaptations to tolerate the lipotoxic events triggered by the elevation in sphingosine and related sphingolipids such as sphingomyelin that interact with cholesterol (Slotte, 1999). This inference is further supported by previous findings that sphingolipid levels were globally elevated in the representative IDH1^wt^ GBM compared to various IDH1^mut^ gliomas subtypes (Dowdy et al., 2020). However, the representative IDH1^wt^ GBM was also less susceptible to the Sph C17 and NDMS combination treatment than IDH1^mut^ gliomas (figure 1c).

Based on the elevated protein expression of ABCA1 in representative IDH1^wt^ GBM compared to various IDH1^mut^ gliomas, it was inferred that ABCA1 was an integral component in the mechanism to protect neurospheres from the lipotoxic effects of treatment. Thus, we examined the drug dose response for IDH1^mut^ neurospheres modified with either of two different ΔABCA1 (900 or 901) knock-out (KO) compared to vector (EV) (figure 2h) over 24 h period. Surprisingly, the outcome conflicted with our original expectation that the IDH1^wt^ GBM benefited from the overexpression of ABCA1 to promote efflux and mitigate lipotoxic effects of accumulation of unesterified cholesterol or sphingolipid substrates. Therefore, we conducted a follow-on analysis to probe whether rescue benefits were encountered for representative IDH1^mut^ neurospheres after being pre-incubated at various doses (0, 50, 250, and 800 nM) of cholesterol (figure 2h) for 2 h before treatment—based on prior optimization to examine the kinetic uptake of cholesterol-NBD within the representative IDH1^mut^ glioma. Surprisingly, the outcome did not yield any rescue benefit that was evident for representative IDH1^mut^ neurospheres following the 24 h combination treatment. Given that the representative IDH1^wt^ GBM was shown to present a characteristically higher level of sphingolipids (e.g., S1P, sphingosine, sphingomyelin) compared to IDH1^mut^ glioma subtypes (Dowdy et al., 2020),it was inferred that IDH1^wt^ GBM must benefit from a unique array of additional mechanistic protections that limit their vulnerability to the lipotoxic effects sphingosine and NDMS accumulation (Dowdy et al., 2020).

Other factors are involved that provide additional insight into the mechanism of action for the drug, particularly in IDH1^mut^ glioma response to the drug. Moreover, the data revealed that expression of LXRα (NR1H3) remained elevated across a range of IDH1^mut^ glioma subtypes ((figure 2j) in response to the Combo treatment and subsequential accumulation of endogenous sphingosines and sphingosine-derivatives (figure 1h-i). In relation, the gene expression of LXR-mediated MYLIP remained elevated across IDH1^mut^ glioma subtypes after 48 h (figure 2k) in accordance with early onset MYLIP expression (figure 2a). This is consistent with the decline in intracellular cholesterol given that MYLIP was previously reported to indirectly suppress the uptake of cholesterol by LDL-receptor (LDLR), the target of MYLIP-driven proteolytic ubiquitination (Adi et al., 2019; Zelcer et al., 2009).

In addition, the Kaplan-Meier survival curve comparison (figure 2l) generated from TGCA revealed that elevated gene expression for NR1H3 may be connected to decreased survivability in LGG clinical patients. This correlation provided additional insight into the protective role of LXR-mediated activity, which may have been altered by drug-related elevation of sphingosines (figure d-i). As expected for LXRα (NR1H3) activation and signaling, the decline in the transcription of HMGCS1 (figure 2m) and HMGCR (figure 2n) was preserved across multiple IDH1^mut^ glioma subtypes in response to combo treatment even after 48 h. Likewise, this trend was reproduced at the protein levels for HMGCS1 and HMGCR1 (figure 2o), which control the rate-limiting onset for the biosynthesis of cholesterol and its precursors (Chen et al., 2022; DeBose-Boyd, 2008; H. yan Wang et al., 2022; Yeh et al., 2018). In addition, the decline in LDLR protein levels (figure 2o) was consistent with elevated transcription for MYLIP (figure 2a and 2k) observed across IDH1^mut^ gliomas subtypes in treatment.

We also observed a decline in the mRNA levels of NPC2 (figure 2p); which is known to promote the export of unesterified cholesterol derived from cholesteryl esters (CEs) from within the lysosome (Heybrock et al., 2019; Storch et al., 2009). Acetyl-CoA acetyltransferase 2 (ACAT2) expression is decreased (figure 2q) despite its role as a cholesterol biosensor that facilitates the esterification of cholesterol to form CEs and lipid droplets (LD) (Storch & Xu, 2009). Typically, the formation of CE is expected to mitigate cholesterol-associated membrane stress in the ER (Hager et al., 2012; Storch & Xu, 2009). The observed elevation in the transcription for LXR-mediated lysophosphatidylcholine acyltransferase 3 (LPCAT3) (figure 2r) was previously associated with suppressing cholesterol biosynthesis to help mitigate stress in ER and suppress tumorigenesis (Shao et al., 2022; B. Wang et al., 2018) Given the decline in ER-associated ACAT2 expression, it is unclear whether the decrease in endo-liposomal NPC2 expression contributed to the evident elevation of certain CEs (Figure 1c-f) in the lipidome. Notably, ACAT2 suppression was previously correlated to elevated ABCA1 expression (Pedrelli et al., 2014).

Both, the early onset (24 h) and late-onset (48 h) expression events provided the valuable insight necessary to shape our understanding of the mechanisms of drug action across IDH1mut glioma subtypes. We interpreted the above correlations as apparent indicators that the drug-dependent activation of LXR-related events triggered a decline in intracellular cholesterol below basal levels (figure 1j-k). It appeared that a critical point was reached, leading to a loss in cholesterol-associated benefits to proliferation and, potentially, membrane integrity. Given that early onset elevation in the transcription of specific indicators of membrane degradation (ROCK1) and the potential for sphingosine to directly trigger nuclear LXR-associated activity, the multi-omics analysis was complemented with confocal fluorescent microscopy to probe the trafficking of fluorescent-labeled cholesterol, sphingosine, or both with and without the drug combination (further explored in figure 3).

**Figure 3.**
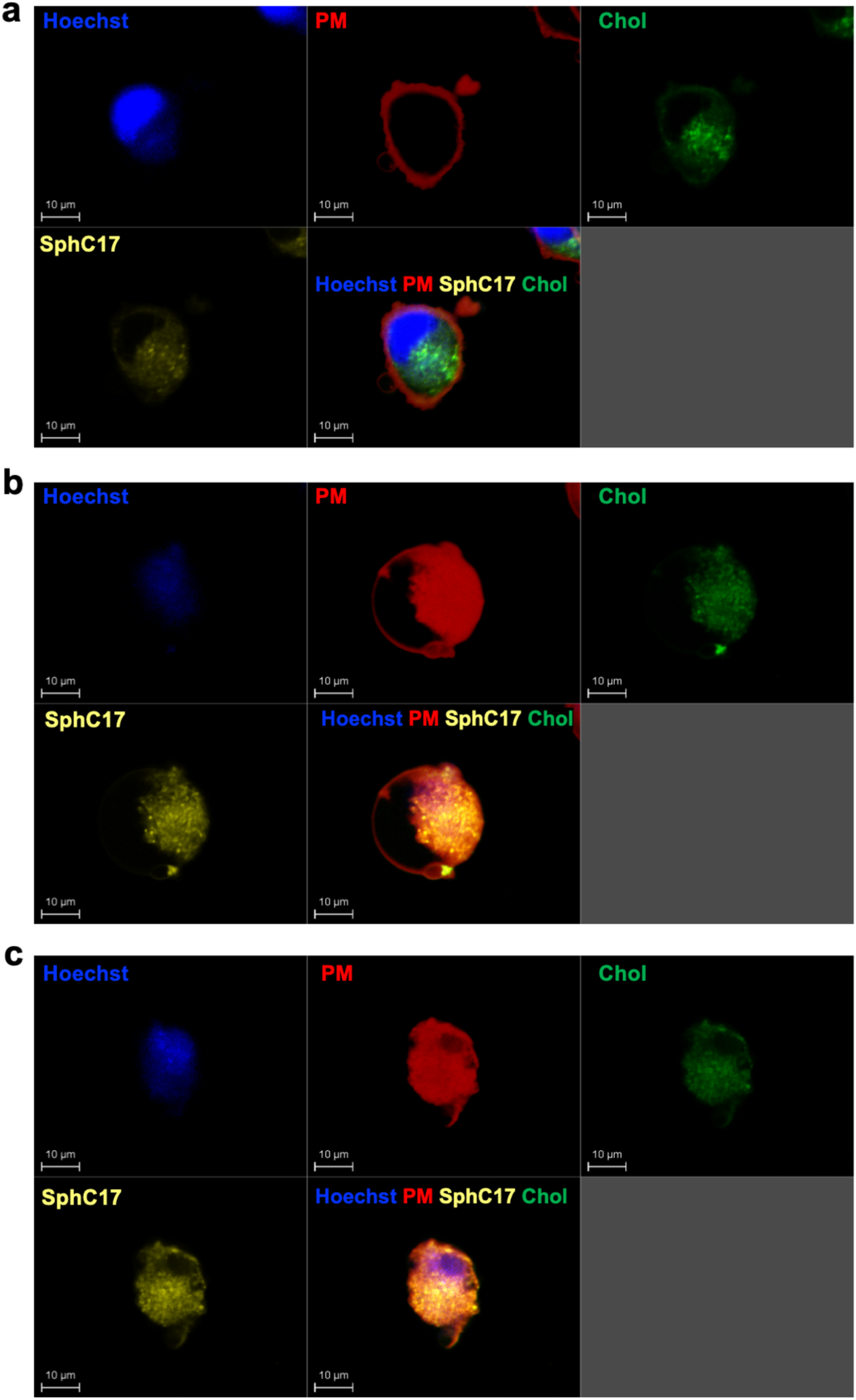
Probing intracellular cholesterol and sphingosine trafficking in IDH1^mut^ glioma with treatment using live cell imaging. Tracking of BODIPY-cholesterol (green) and Rhodamine C17 Sphingosine (yellow) reveals that they accumulate at the plasma membrane (red). Concurrent staining of the nucleus (blue) reveals that cytosolic cholesterol is trafficked to the nucleus, where it becomes concentrated and causes the blebbing of the membrane. a: frame 1 (t∼30 min since the addition of the combo); b: frame 10 (t=130 min c: frame 20 (t=230 min).

### Sphingosine C17 addition recruits’ cholesterol at the plasma membrane

To understand the mechanism by which sphingosine alters cholesterol homeostasis and leads to such a drastic decrease in cholesterol levels inside the cells, we did live cell confocal microscopy. When we incubated the cells 500 nM sphingosine C17 (Rhodamine) and 500 nM BODIPY-Cholesterol for 24 hours followed but washing and staining with Hoechst (blue) and plasma membrane(red) (Fig 3a-c). Within two hours of combo addition, the membrane starts to sweal, leading to blebbing.

## DISCUSSION

While SphC17 was administered to promote the elevation of intracellular sphingosine levels, NDMS is a potent SphK1 inhibitor that was also administered to suppress the ability of IDH1^mut^ gliomas to mitigate the overwhelming accumulation of sphingosine via production to pro-oncogenic S1P C18. In turn, the LC/MS analysis revealed that the primary substrate for SphK, endogenous sphingosine C18, was among the most significant sphingolipids to be profoundly elevated across representative IDH1^mut^ (oligodendroglioma, astrocytoma, and U251-IDH1^R132H^ overexpression model) neurospheres in a dose-dependent fashion. Although a relative decline in S1P was observed, the scale was unremarkable. Based on these findings, it was inferred that the inhibition of SphK activity was unlikely the primary mechanism of drug action. The significant elevation of sphingosine C16, C17, and C18 was particularly interesting, given that prior studies reported that the elevation of sphingosine can trigger apoptosis through a mechanism like the accumulation of ceramide (e.g., ER-stress) (Dowdy et al., 2020; Taha et al., 2006). In contrast, differential expression of ER-stress associated markers was not observed consistently. However, the results of the immunoblot analysis revealed that apoptotic markers (Parp1α and CytC) consistent with mitochondria-stress induced apoptosis increased over time in response to drug treatment.

We report a global downregulation in the transcription for the gateway enzymes along with all other enzymes that participate in biosynthesis and cholesterol, sterols, and precursor isoprenoids within the treated IDH1^mut^ gliomas. These trends provide evidence that the sphingosine-based treatment activates LXRα (NR1H3) signaling. LXR signaling can modulate cholesterol homeostasis by activating expression genes, ABCA1 and ABCG1 involved in reverse cholesterol transport; MYLIP that activate ubiquitinated digestion of influx protein LDLR; and suppression of the cascade genes involved in cholesterol and precursor biosynthesis (Rong et al., 2013; Tarling et al., 2011; Yvan-Charvet et al., 2010; Zelcer et al., 2009; Zhu et al., 2012). Complementary to transcriptomic RNA expression data, immunoblot analysis also demonstrated a decline for enzymes involved (HMGCR and HMGCS) in the rate-limiting entry for cholesterol, sterol, and isoprenoid biosynthesis(Chen et al., 2022; DeBose-Boyd, 2008; Yeh et al., 2018).

While the gene expression of MYLIP (which is typically modulated by LXRα (Xu et al., 2018; Zelcer et al., 2009)was elevated, the immunoblot data revealed that the related LDL-receptor (LDLR) protein was decreased in response to drug treatment. While plasma membrane LDLR facilitates the docking of LDL-cholesterol and mediates internalization via endocytosis, LDLR protein levels can be negatively impacted by membrane cholesterol levels and polyubiquitinated degradation driven by LXRα-dependent MYLIP activity (Morales et al., 2023). Thus, MYLIP-driven degradation of LDLR protein could hinder cholesterol uptake and internalization.

Several targets of LXRα signaling (NR1H3, ABCA1, ABCG1, MYLIP, and LPCAT3) were significantly upregulated following drug treatment (highlighted in the volcano plot). Furthermore, the global expression of genes directly involved in cholesterol biosynthesis (figure 2a) was among the most significantly suppressed genes in the transcriptome. The LXRα-driven suppression of genes involved in the cholesterol biosynthesis pathway can be triggered by the accumulation of cytotoxic cholesterol levels (Morales et al., 2023; Widenmaier et al., 2017; Zhao et al., 2010). This may serve as a rescue response to mitigate ER stress due to the saturation of membrane cholesterol.

An additional understanding of the connection between the elevation of sphingosine and cholesterol was acquired through the application of fluorescent microscopy. While tracking the cholesterol using fluorescent probes, it was revealed that cholesterol that readily permeates the plasma membrane accumulates in the lysosome, ER, and eventually encompasses the nucleus, where nuclear receptors can actively regulate cholesterol homeostasis. Follow-on fluorescent probing of sphingosine revealed that sphingosine trafficking mimics the behavior of cholesterol eventually accumulating at the nuclear membrane. Upon combining the cholesterol and sphingosine-containing fluorophores, a cohesive relationship was observed involving concerted trafficking to and from the nucleus.

Based on microscopy with a representative fluorophore, the observation that a cohesive interaction occurs between unesterified cholesterol and specific sphingolipids (e.g., sphingosine) when elevated is also consistent with previous research (Altuzar et al., 2023; Frisz et al., 2013; Garmy et al., 2005; McIntosh et al., 1992; Samsonov et al., 2001; Zhang et al., 2019). The combined accumulation of these molecules at the nucleus triggered their concerted trafficking of sphingosine and associated cholesterol to saturate the PM within the initial 8h. Follow-on conformation changes ensued across the outer plasma membrane. Despite the known protective properties of cholesterol (Biswas et al., 2019), the saturation of the PM with sphingosine-associated cholesterol resulted in swelling, vesiculation, and progressive membrane degradation that resembles pyroptotic programmed cell death (Gong et al., 2020). Following 24 h treatment with SphC17 and NDMS combination, the occurrence of swelling and blebbing of the PM was also consistent with the early elevation of specific markers, ROCK1 and ROCK2, (Figure 2a) involved apoptotic membrane blebbing and vesiculation (Aoki et al., 2020; Coleman et al., 2001; Leverrier & Ridley, 2001; Sebbagh et al., 2001). Labeling of plasma membrane further validates the blebbing via live cell microscopy (Fig 3).

This multi-omics investigation on the mechanism of drug action has elucidated a remarkable, interactive association between sphingosine and cholesterol consistent with early research (Altuzar et al., 2023; Garmy et al., 2005). The elevation of sphingosine and NDMS (sphingosine-derived SphK inhibitor) disrupted the expression of gene and catalytic enzymes involved in the cholesterol metabolism, while promoting the gene and protein expression for crucial components of the cholesterol efflux, ABCA1 and ABCG1. While these observed responses are typically part of LXRα-driven rescue from excess accumulation of cytotoxic cholesterol, LC/MS analysis determined that cholesterol levels were significantly declined and not elevated over the time-course between 0.5 h to 24 h following drug treatment.

Follow-on fluorescent probing with sphingosine and cholesterol provided visible evidence that the elevation of sphingosine triggers their cohesive migration and saturation in the plasma membrane and subsequent vesiculation. Interestingly, apoptotic membrane blebbing leads to the release of vesicles from the PM, which is considered a profound indicator of membrane disintegration (Aoki et al., 2020; Leverrier & Ridley, 2001; Lima et al., 2017). Recent research has discovered new ways of programmed cell death known as pyroptosis results in membrane perforation, cell swelling, membrane blebbing is, and cell lysis (Gong et al., 2020). Independent of caspase 3, pyroptosis relies on caspases (caspase-1 and caspase-4/5/11) that are accompanied by inflammation (Gong et al., 2020).

## CONCLUSIONS

Overall, these findings provide insight into complex interactions that are mediated by the balancing of sphingolipid metabolism, as previously proposed by Dowdy et al. (2020). Based on previous research (Dowdy et al., 2020), the potent biostatic effect of the drug on highly susceptible IDH1^mut^ oligodendrogliomas and astrocytomas (Dowdy et al., 2020) was initially hypothesized to result from a disruption of the rheostatic balance between S1P, sphingosine, and ceramide. This investigation revealed a unique, remarkable correlation between sphingosine and cholesterol underlying the mechanism of action for the proposed combination treatment. While it is well known that activation of LXRα nuclear receptor signaling serves a key role in balancing intracellular cholesterol homeostasis, LXRα can promote the expression of genes and related proteins that promote efflux of cholesterol (ABCA1 and ABCG1), regulate influx while suppressing the expression of genes involved in cholesterol biosynthesis pathways. In this study, glioma neurospheres presented upregulated gene and protein expression for ABCA1 and ABCG1 transporters for cholesterol efflux and complementary downregulation gene and protein expression related to cholesterol biosynthesis in a drug dose- and time-dependent fashion.

Based on the evidence from the multi-omics investigation, we determined that the proposed drug treatment led to a direct, global elevation of sphingosine along with other vital sphingolipids (i.e., SM and ceramide). In turn, the decline in intracellular cholesterol was primarily facilitated by the activation of cholesterol homeostasis signaling pathways, leading to the upregulation of genes involving cholesterol efflux and suppression of genes involving cholesterol biosynthesis. Given that the signaling of cholesterol regulation was unlikely driven by a cytotoxic elevation of free cholesterol, we propose that saturation of the cellular membranes with unesterified cholesterol in association with global elevation in sphingosine could account for the activation of the cholesterol efflux and the suppression of cholesterol biosynthesis (Figure 4).

**Figure 4.**
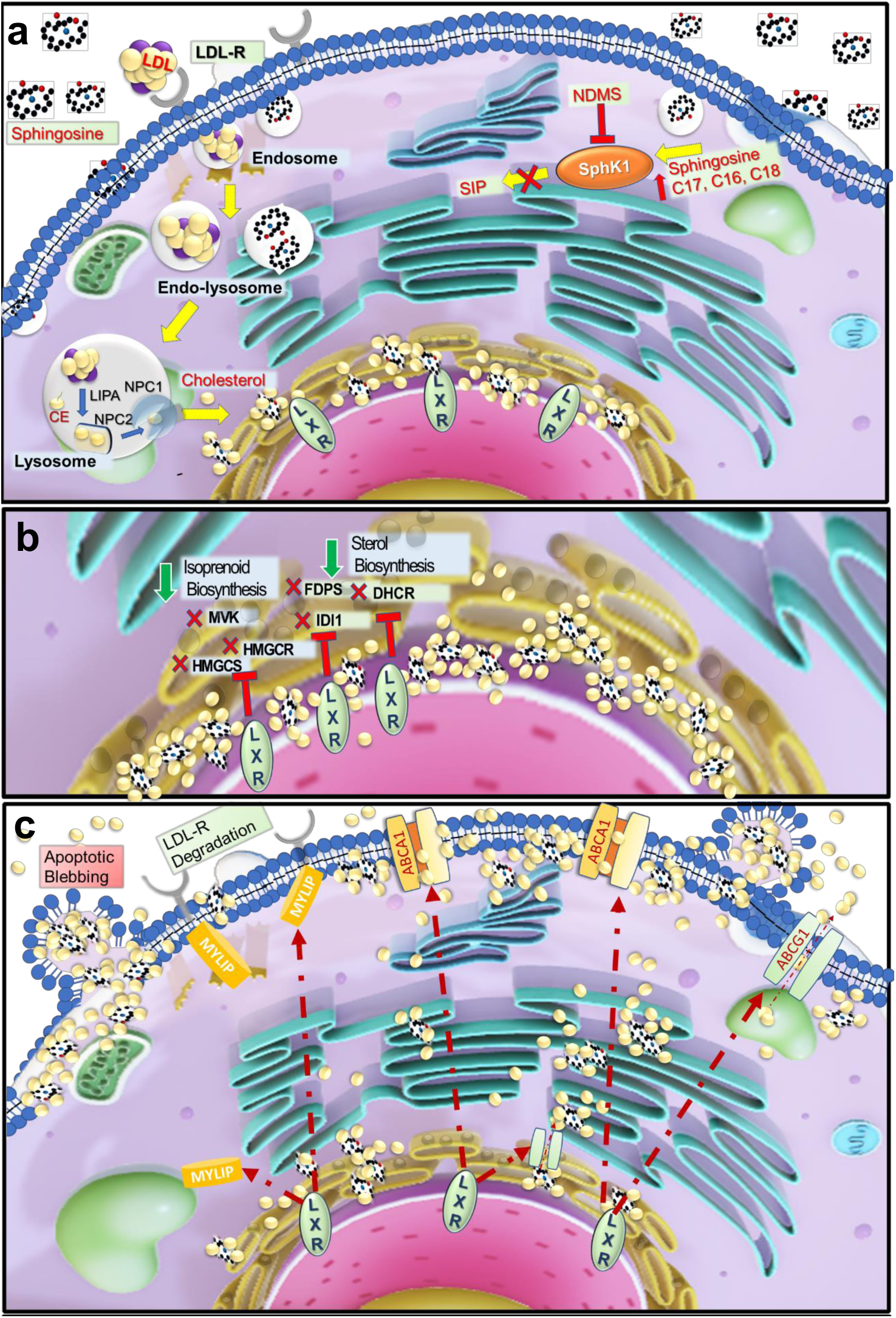
Sphingosine addition recruits cholesterol to the plasma membrane blebbing and apoptosis. a. Early events of sphingosine addition and cholesterol. a. Cholesterol accumulation triggers, LXR signaling and a decrease of sterol synthesis. c. Cholesterol is sequestered at the plasma membrane by sphingosine leading to membrane blebbing and apoptosis.

This proposed rescue response provided further insight into the mechanism of drug action which led to the membrane saturation with sphingosine and cholesterol. We concluded that the saturation of the plasma membrane with sphingosines and associated cholesterol triggered the plasma membrane degradation resembling pyroptosis (Fig. 3b). The unique correlation between the elevation of sphingosine (and sphingosine derivative, NMDS) and the regulation of intracellular cholesterol, as described herein, offers insight into how the dysregulation of certain sphingolipids can modulate signaling. The novel revelation that reprogramming sphingolipid metabolism can effectively arrest the proliferation of IDH1^mut^ glioma offers a direction for the advancements of novel treatments and warrants ongoing investigation as a preclinical model and potential therapeutic intervention to improve patient outcomes.

## Author Contributions

Conceptualization, T.D. and M.L.; methodology, T.D., H.M. A.L., and M.L; software, T.D., A.L., A.L., and M.L.; validation, T.D., M.L., A.L. H.M, T.Y.; formal analysis, T.D, and M.L.; data curation, T.D.; writing—original draft preparation, T.D.; writing—review and editing, T.D., M.L., A.L., A.L. L.Z., T.Y., M.G., and M.L.; supervision, M.L.; project administration, M.L.; funding acquisition, M.G. All authors have read and agreed to the published version of the manuscript.

## Acknowledgments

The authors would like to thank Alan and Ashley Dabbiere for their monetary donation which contributed to the acquisition of the Leica Stellaris 8 CRS instrument.

**Acknowledgments:** We would like to thank Dr. ChunZhang Yang (NOB) for providing us with the U251*^WT^*and U251 *^R132H^* constructs and Dr. Timothy Chan for providing us with TS603 cells.

## References

Adi, D., Lu, X. Y., Fu, Z. Y., Wei, J., Baituola, G., Meng, Y. J., Zhou, Y. X., Hu, A., Wang, J. K., Lu, X. F., Wang, Y., Song, B. L., Ma, Y. T., & Luo, J. (2019). IDOL G51S Variant Is Associated with High Blood Cholesterol and Increases Low-Density Lipoprotein Receptor Degradation. *Arteriosclerosis*, Thrombosis, and Vascular Biology, 39(12). 10.1161/ATVBAHA.119.312589

Alnaaim, S. A., Al-Kuraishy, H. M., Alexiou, A., Papadakis, M., Saad, H. M., & Batiha, G. E. S. (2024). Role of Brain Liver X Receptor in Parkinson’s Disease: Hidden Treasure and Emerging Opportunities. In Molecular Neurobiology (Vol. 61, Issue 1). 10.1007/s12035-023-03561-y

Altuzar, J., Notbohm, J., Stein, F., Haberkant, P., Hempelmann, P., Heybrock, S., Worsch, J., Saftig, P., & Höglinger, D. (2023). Lysosome-targeted multifunctional lipid probes reveal the sterol transporter NPC1 as a sphingosine interactor. Proceedings of the National Academy of Sciences of the United States of America, 120(11). 10.1073/pnas.2213886120

Aoki, K., Satoi, S., Harada, S., Uchida, S., Iwasa, Y., & Ikenouchi, J. (2020). Coordinated changes in cell membrane and cytoplasm during maturation of apoptotic bleb. Molecular Biology of the Cell, 31(8). 10.1091/MBC.E19-12-0691

Appelqvist, H., Nilsson, C., Garner, B., Brown, A. J., Kågedal, K., & Öllinger, K. (2011). Attenuation of the lysosomal death pathway by lysosomal cholesterol accumulation. American Journal of Pathology, 178(2). 10.1016/j.ajpath.2010.10.030

Biswas, A., Kashyap, P., Datta, S., Sengupta, T., & Sinha, B. (2019). Cholesterol Depletion by MβCD Enhances Cell Membrane Tension and Its Variations-Reducing Integrity. Biophysical Journal, 116(8). 10.1016/j.bpj.2019.03.016

Celiku, O., Johnson, S., Zhao, S., Camphausen, K., & Shankavaram, U. (2014). Visualizing molecular profiles of glioblastoma with GBM-BioDP. PLoS ONE, 9(7). 10.1371/journal.pone.0101239

Chen, Y., Li, M., Yang, Y., Lu, Y., & Li, X. (2022). Antidiabetic drug metformin suppresses tumorigenesis through inhibition of mevalonate pathway enzyme HMGCS1. Journal of Biological Chemistry, 298(12). 10.1016/j.jbc.2022.102678

Clouser-Roche, A., Johnson, K., Fast, D., & Tang, D. (2008). Beyond pass/fail: A procedure for evaluating the effect of carryover in bioanalytical LC/MS/MS methods. Journal of Pharmaceutical and Biomedical Analysis, 47(1). 10.1016/j.jpba.2007.12.019

Coleman, M. L., Sahai, E. A., Yeo, M., Bosch, M., Dewar, A., & Olson, M. F. (2001). Membrane blebbing during apoptosis results from caspase-mediated activation of ROCK I. Nature Cell Biology, 3(4). 10.1038/35070009

DeBose-Boyd, R. A. (2008). Feedback regulation of cholesterol synthesis: Sterol-accelerated ubiquitination and degradation of HMG CoA reductase. Cell Research, 18(6). 10.1038/cr.2008.61

Dowdy, T., Zhang, L., Celiku, O., Movva, S., Lita, A., Ruiz-Rodado, V., Gilbert, M. R., & Larion, M. (2020). Sphingolipid pathway as a source of vulnerability in IDH1mut glioma. Cancers, 12(10). 10.3390/cancers12102910

Feramisco, J. D., Goldstein, J. L., & Brown, M. S. (2004). Membrane Topology of Human Insig-1, a Protein Regulator of Lipid Synthesis. Journal of Biological Chemistry, 279(9). 10.1074/jbc.M312623200

Frisz, J. F., Lou, K., Klitzing, H. A., Hanafin, W. P., Lizunov, V., Wilson, R. L., Carpenter, K. J., Kim, R., Hutcheon, I. D., Zimmerberg, J., Weber, P. K., & Kraft, M. L. (2013). Direct chemical evidence for sphingolipid domains in the plasma membranes of fibroblasts. Proceedings of the National Academy of Sciences of the United States of America, 110(8). 10.1073/pnas.1216585110

Garmy, N., Taïeb, N., Yahi, N., & Fantini, J. (2005). Interaction of cholesterol with sphingosine. Journal of Lipid Research, 46(1). 10.1194/jlr.m400199-jlr200

Gong, W., Shi, Y., & Ren, J. (2020). Research progresses of molecular mechanism of pyroptosis and its related diseases. In Immunobiology (Vol. 225, Issue 2). 10.1016/j.imbio.2019.11.019

Hager, L., Li, L., Pun, H., Liu, L., Hossain, M. A., Maguire, G. F., Naples, M., Baker, C., Magomedova, L., Tam, J., Adeli, K., Cummins, C. L., Connelly, P. W., & Ng, D. S. (2012). Lecithin:cholesterol acyltransferase deficiency protects against cholesterol-induced hepatic endoplasmic reticulum stress in mice. Journal of Biological Chemistry, 287(24). 10.1074/jbc.M112.340919

Heybrock, S., Kanerva, K., Meng, Y., Ing, C., Liang, A., Xiong, Z. J., Weng, X., Ah Kim, Y., Collins, R., Trimble, W., Pomès, R., Privé, G. G., Annaert, W., Schwake, M., Heeren, J., Lüllmann-Rauch, R., Grinstein, S., Ikonen, E., Saftig, P., & Neculai, D. (2019). Lysosomal integral membrane protein-2 (LIMP-2/SCARB2) is involved in lysosomal cholesterol export. Nature Communications, 10(1). 10.1038/s41467-019-11425-0

Huotari, J., & Helenius, A. (2011). Endosome maturation. In EMBO Journal (Vol. 30, Issue 17). 10.1038/emboj.2011.286

Jiang, X. C., & Li, Z. (2022). Sphingolipids and Cholesterol. In Advances in Experimental Medicine and Biology (Vol. 1372). 10.1007/978-981-19-0394-6_1

Larion, M., Dowdy, T., Ruiz-Rodado, V., Meyer, M. W., Song, H., Zhang, W., Davis, D., Gilbert, M. R., & Lita, A. (2019). Detection of metabolic changes induced via drug treatments in live cancer cells and tissue using raman imaging microscopy. Biosensors, 9(1). 10.3390/bios9010005

Leverrier, Y., & Ridley, A. J. (2001). Apoptosis: Caspases orchestrate the ROCK “n” bleb. Nature Cell Biology, 3(4). 10.1038/35070151

Lima, S., Milstien, S., & Spiegel, S. (2017). Sphingosine and sphingosine kinase 1 involvement in endocytic membrane trafficking. Journal of Biological Chemistry, 292(8). 10.1074/jbc.M116.762377

McIntosh, T. J., Simon, S. A., Needham, D., & Huang, C. hsien. (1992). Structure and Cohesive Properties of Sphingomyelin/Cholesterol Bilayers. Biochemistry, 31(7). 10.1021/bi00122a017

Morales, S. V., Mahmood, A., Pollard, J., Mayne, J., Figeys, D., & Wiseman, P. W. (2023). The LDL receptor is regulated by membrane cholesterol as revealed by fluorescence fluctuation analysis. Biophysical Journal, 122(18). 10.1016/j.bpj.2023.08.005

Pang, Z., Chong, J., Zhou, G., De Lima Morais, D. A., Chang, L., Barrette, M., Gauthier, C., Jacques, P. É., Li, S., & Xia, J. (2021). MetaboAnalyst 5.0: Narrowing the gap between raw spectra and functional insights. Nucleic Acids Research, 49(W1). 10.1093/nar/gkab382

Pedrelli, M., Davoodpour, P., Degirolamo, C., Gomaraschi, M., Graham, M., Ossoli, A., Larsson, L., Calabresi, L., Gustafsson, J. Å., Steffensen, K. R., Eriksson, M., & Parini, P. (2014). Hepatic ACAT2 knock down increases ABCA1 and modifies HDL metabolism in mice. PLoS ONE, 9(4). 10.1371/journal.pone.0093552

Phillips, R. E., Yang, Y., Smith, R. C., Thompson, B. M., Yamasaki, T., Soto-Feliciano, Y. M., Funato, K., Liang, Y., Garcia-Bermudez, J., Wang, X., Garcia, B. A., Yamasaki, K., McDonald, J. G., Birsoy, K., Tabar, V., & Allis, C. D. (2019). Target identification reveals lanosterol synthase as a vulnerability in glioma. Proceedings of the National Academy of Sciences of the United States of America, 116(16). 10.1073/pnas.1820989116

Qiu, Z., Yuan, W., Chen, T., Zhou, C., Liu, C., Huang, Y., Han, D., & Huang, Q. (2016). HMGCR positively regulated the growth and migration of glioblastoma cells. Gene, 576(1). 10.1016/j.gene.2015.09.067

Röhrl, C., Eigner, K., Winter, K., Korbelius, M., Obrowsky, S., Kratky, D., Kovacs, W. J., & Stangl, H. (2014). Endoplasmic reticulum stress impairs cholesterol efflux and synthesis in hepatic cells. Journal of Lipid Research, 55(1). 10.1194/jlr.M043299

Rong, X., Albert, C. J., Hong, C., Duerr, M. A., Chamberlain, B. T., Tarling, E. J., Ito, A., Gao, J., Wang, B., Edwards, P. A., Jung, M. E., Ford, D. A., & Tontonoz, P. (2013). LXRs regulate ER stress and inflammation through dynamic modulation of membrane phospholipid composition. Cell Metabolism, 18(5). 10.1016/j.cmet.2013.10.002

Samsonov, A. V., Mihalyov, I., & Cohen, F. S. (2001). Characterization of cholesterol-sphingomyelin domains and their dynamics in bilayer membranes. Biophysical Journal, 81(3). 10.1016/S0006-3495(01)75803-1

Sebbagh, M., Renvoizé, C., Hamelin, J., Riché, N., Bertoglio, J., & Bréard, J. (2001). Caspase-3-mediated cleavage of ROCK I induces MLC phosphorylation and apoptotic membrane blebbing. Nature Cell Biology, 3(4). 10.1038/35070019

Sever, N., Yang, T., Brown, M. S., Goldstein, J. L., & DeBose-Boyd, R. A. (2003). Accelerated degradation of HMG CoA reductase mediated by binding of insig-1 to its sterol-sensing domain. Molecular Cell, 11(1). 10.1016/S1097-2765(02)00822-5

Shao, G., Qian, Y., Lu, L., Liu, Y., Wu, T., Ji, G., & Xu, H. (2022). Research progress in the role and mechanism of LPCAT3 in metabolic related diseases and cancer. In Journal of Cancer (Vol. 19, Issue 8). 10.7150/jca.71619

Slotte, J. P. (1999). Sphingomyelin - Cholesterol interactions in biological and model membranes. Chemistry and Physics of Lipids, 102(1–2). 10.1016/S0009-3084(99)00071-7

Storch, J., & Xu, Z. (2009). Niemann-Pick C2 (NPC2) and intracellular cholesterol trafficking. In Biochimica et Biophysica Acta - Molecular and Cell Biology of Lipids (Vol. 1791, Issue 7). 10.1016/j.bbalip.2009.02.001

Taha, T. A., Mullen, T. D., & Obeid, L. M. (2006). A house divided: Ceramide, sphingosine, and sphingosine-1-phosphate in programmed cell death. In Biochimica et Biophysica Acta - Biomembranes (Vol. 1758, Issue 12). 10.1016/j.bbamem.2006.10.018

Tarling, E. J., & Edwards, P. A. (2011). ATP binding cassette transporter G1 (ABCG1) is an intracellular sterol transporter. Proceedings of the National Academy of Sciences of the United States of America, 108(49). 10.1073/pnas.1113021108

Tixeira, R., Phan, T. K., Caruso, S., Shi, B., Atkin-Smith, G. K., Nedeva, C., Chow, J. D. Y., Puthalakath, H., Hulett, M. D., Herold, M. J., & Poon, I. K. H. (2020). ROCK1 but not LIMK1 or PAK2 is a key regulator of apoptotic membrane blebbing and cell disassembly. Cell Death and Differentiation, 27(1). 10.1038/s41418-019-0342-5

Trinh, M. N., Brown, M. S., Goldstein, J. L., Han, J., Vale, G., McDonald, J. G., Seemann, J., Mendell, J. T., & Lu, F. (2020). Last step in the path of LDL cholesterol from lysosome to plasma membrane to ER is governed by phosphatidylserine. Proceedings of the National Academy of Sciences of the United States of America, 117(31). 10.1073/pnas.2010682117

Wang, B., Rong, X., Palladino, E. N. D., Wang, J., Fogelman, A. M., Martín, M. G., Alrefai, W. A., Ford, D. A., & Tontonoz, P. (2018). Phospholipid Remodeling and Cholesterol Availability Regulate Intestinal Stemness and Tumorigenesis. Cell Stem Cell, 22(2). 10.1016/j.stem.2017.12.017

Wang, H. yan, Yu, P., Chen, X. sha, Wei, H., Cao, S. jie, Zhang, M., Zhang, Y., Tao, Y. guang, Cao, D. sheng, Qiu, F., & Cheng, Y. (2022). Identification of HMGCR as the anticancer target of physapubenolide against melanoma cells by in silico target prediction. Acta Pharmacologica Sinica, 43(6). 10.1038/s41401-021-00745-x

Wang, T., Zhao, Y., You, Z., Li, X., Xiong, M., Li, H., & Yan, N. (2020). Endoplasmic reticulum stress affects cholesterol homeostasis by inhibiting LXRα expression in hepatocytes and macrophages. Nutrients, 12(10). 10.3390/nu12103088

Widenmaier, S. B., Snyder, N. A., Nguyen, T. B., Arduini, A., Lee, G. Y., Arruda, A. P., Saksi, J., Bartelt, A., & Hotamisligil, G. S. (2017). NRF1 Is an ER Membrane Sensor that Is Central to Cholesterol Homeostasis. Cell, 171(5). 10.1016/j.cell.2017.10.003

Xu, Y., Hernández-Ledezma, J. J., Hutchison, S. M., & Bogan, R. L. (2018). The liver X receptors and sterol regulatory element binding proteins alter progesterone secretion and are regulated by human chorionic gonadotropin in human luteinized granulosa cells. Molecular and Cellular Endocrinology, 473. 10.1016/j.mce.2018.01.011

Yeh, Y. S., Jheng, H. F., Iwase, M., Kim, M., Mohri, S., Kwon, J., Kawarasaki, S., Li, Y., Takahashi, H., Ara, T., Nomura, W., Kawada, T., & Goto, T. (2018). The Mevalonate Pathway Is Indispensable for Adipocyte Survival. IScience, 9. 10.1016/j.isci.2018.10.019

Yvan-Charvet, L., Pagler, T. A., Seimon, T. A., Thorp, E., Welch, C. L., Witztum, J. L., Tabas, I., & Tall, A. R. (2010). ABCA1 and ABCG1 protect against oxidative stress-induced macrophage apoptosis during efferocytosis. Circulation Research, 106(12). 10.1161/CIRCRESAHA.110.217281

Zaibaq, F., Dowdy, T., & Larion, M. (2022). Targeting the Sphingolipid Rheostat in Gliomas. In International Journal of Molecular Sciences (Vol. 23, Issue 16). 10.3390/ijms23169255

Zelcer, N., Hong, C., Boyadjian, R., & Tontonoz, P. (2009). LXR regulates cholesterol uptake through idol-dependent ubiquitination of the LDL receptor. Science, 325(5936). 10.1126/science.1168974

Zhang, Y., Zhang, J., Li, Q., Wu, Y., Wang, D., Xu, L., Zhang, Y., Wang, S., Wang, T., Liu, F., Zaky, M. Y., Hou, S., Liu, S., Zou, K., Lei, H., Zou, L., & Liu, H. (2019). Cholesterol content in cell membrane maintains surface levels of ErbB2 and confers a therapeutic vulnerability in ErbB2-positive breast cancer. Cell Communication and Signaling, 17(1). 10.1186/s12964-019-0328-4

Zhao, C., & Dahlman-Wright, K. (2010). Liver X receptor in cholesterol metabolism. In Journal of Endocrinology (Vol. 204, Issue 3). 10.1677/JOE-09-0271

Zhu, R., Ou, Z., Ruan, X., & Gong, J. (2012). Role of liver X receptors in cholesterol efflux and inflammatory signaling (review). In Molecular Medicine Reports (Vol. 5, Issue 4). 10.3892/mmr.2012.758

